# PRMT7 restricts CD8^+^ T cells expansion via the NF-κB pathway

**DOI:** 10.64898/2026.04.25.720790

**Authors:** Nivine Srour, Kaixiu Luo, David Allard, Yan Xiong, Zhenbao Yu, Theodore Papadopoulos, François Santinon, Cheng-Hsun Hsieh, John Stagg, Dalia Barsyte-Lovejoy, Sonia V del Rincon, Jian Jin, Stéphane Richard

## Abstract

New therapeutics are needed to enhance cytotoxic T lymphocyte (CTL) expansion and effector function for effective tumor control. Here, we show that T cell specific deletion of Prmt7 using CD4-Cre mice increases CD8^+^ effector differentiation, cytokine secretion, cytolytic activity, and anti-tumor responses. Prmt7 deficiency transcriptionally reprogrammed CD8^+^ T cells by activating the NF-κB pathway, boosting proliferation, and elevating effector molecules such as CD25, CD69, and IFNγ. Mechanistically, PRMT7 associated with RelA and restricted its nuclear translocation. To enable therapeutic translation, we developed a PRMT7-targeting PROTAC degrader (MS54). MS54-treated CTLs exhibited NF-κB pathway activation similar to Prmt7-deficient CTLs. Adoptive transfer of MS54-treated OT-I CTLs significantly improved tumor control in a syngeneic melanoma model. In human CTLs, MS54 enhanced proliferation, activation markers (CD69, CD137), IFNγ production, and cytotoxicity toward melanoma. Together, these findings identify PRMT7 as a negative regulator of CD8^+^ T cell immunity and highlight MS54 as a promising strategy to improve adoptive T cell therapy.

## Introduction

Melanoma, a highly aggressive form of skin cancer, rapidly progresses from primary tumor to metastatic disease, with a rising incidence globally (Centeno *et al*, 2023). Therapeutic strategies include surgical intervention, chemotherapy, radiation, and targeted therapies such as MAP kinase inhibitors for *BRAF* mutated melanoma. Immunotherapies, particularly immune checkpoint inhibitors (ICIs) like anti-PD-1 and anti-CTLA-4 (Larkin *et al*, 2019) and adoptive cell therapies (ACTs), such as chimeric antigen receptor (CAR)-T cells, T cell receptor (TCR)-engineered T cells and tumor-infiltrating lymphocyte (TIL) therapy (Chesney *et al*, 2022; Jilani *et al*, 2024), have shown revolutionary survival outcomes in patients with metastatic melanoma. However immune resistance and T cell dysfunction remain major challenges. Therefore, therapies that enhance CD8^+^ T cell expansion, effector function and amplify antitumor immunity, while minimizing toxicity are highly sought after to improve ACT and ICI efficacy.

Protein arginine methyltransferases (PRMTs) are a family of nine enzymes (PRMT1 to 9) that catalyze arginine methylation and regulate diverse biological processes, including gene expression, RNA splicing, and DNA damage responses (Xu & Richard, 2021). Based on their methylation patterns, PRMTs are classified into type I (e.g., PRMT1–4, 6, 8), type II (PRMT5, 9), and type III (PRMT7) (Bedford & Clarke, 2009). PRMT7 is unique as the only type III enzyme, producing solely monomethylarginine (MMA) (Jain & Clarke, 2019). PRMT7 plays important roles in stress responses, innate immunity via double-stranded RNA signaling, and epigenetic regulation of endogenous retroviral elements (ERVs), thereby influencing tumor immunogenicity (Haghandish *et al*, 2019; Srour *et al*, 2022; Yang *et al*, 2024). Few physiological PRMT7 substrates and interacting proteins have been identified and substrates include histones (Feng *et al*, 2013), HSP70 (Szewczyk *et al*, 2020), and MAVS (Yang *et al*., 2024; Zhu *et al*, 2021).

PRMT7 is overexpressed in breast, colorectal, prostate cancer, as well as chronic myeloid leukemia (CML) (Baldwin *et al*, 2015; Liu *et al*, 2022; Yao *et al*, 2014). Although a potent PRMT7 inhibitor (SGC8158) exists, its poor cell permeability required the design of the prodrug form (SGC3027) (Szewczyk *et al*., 2020). However, no PROteolysis TArgeting Chimera (PROTAC) degraders, which promote selective protein degradation by harnessing the ubiquitin-proteasome system, have yet been reported for PRMT7, unlike PROTACs that specifically target PRMT5 and PRMT6 (Shen *et al*, 2020a; Shen *et al*, 2020b).

Herein we show that conditional deletion of Prmt7 in T cells transcriptionally reprograms CD8^+^ T cells, enhancing their proliferative capacity and upregulating effector molecules such as CD25, CD69 and IFNγ enhancing cytotoxic activity towards cancer cells. To potentially translate our results to the clinic, we developed the first-in-class PRMT7 PROTAC degrader (MS54) that depletes PRMT7 in mouse and human T cells. Human CD8^+^ T cells treated with MS54 had increased proliferation as well as enhanced effector markers including CD69 and CD137 and cytotoxic killing activity. Our findings show that MS54 is a potential therapeutic tool to enhance CD8^+^ T cells expansion and effector function. Mechanistically, in T cells, our data show that PRMT7 fulfills a non-catalytic role as an accessory protein for the p65 (RelA) subunit of NF-κB, restricting its nuclear translocation. Our findings show that MS54 is an effective small molecule to enhance the CD8^+^ T cell response for effective adoptive cell therapy for cancer.

## Results

### *Prmt7^FL/FL; CD4-Cre^* (*Prmt7* cKO) mice have increased CD8^+^ T cells in the periphery

Prmt7 is highly expressed in the thymus and in CD8⁺ T cells compared to other immune cell types (Supplemental Figures S1A, S1B). To define the role of Prmt7 in T cells, we crossed our *Prmt7^FL/FL^* mice (Blanc *et al*, 2016) with *CD4-Cre* transgenic mice in a C57BL/6J background to generate *Prmt7^FL/FL; CD4-Cre^* (*Prmt7* cKO) mice. The mice were healthy with no signs of being immunodeficient (data not shown). Thymocyte extracts were immunoblotted with anti-PRMT7 antibodies and confirmed the absence of Prmt7 in the *Prmt7* cKO mice (Figure 1A). Thymic T cell subpopulations were examined and the loss of Prmt7 did not alter the CD4 and/or CD8 populations (Figures 1B, 1C), suggesting that Prmt7 does not regulate T cell development. Next, we isolated splenocytes from *Prmt7^FL/FL^* (control) and *Prmt7* cKO mice and we observed an increase of ∼ 10% of total CD3^+^ T cells in the *Prmt7* cKO mice compared to control mice (Figures 1D, 1E). We observed a higher CD4^+^CD8^+^ double positive T cells and CD4^-^CD8^+^ single positive T cells, but not CD4^+^ CD8^−^ single positive T cells in *Prmt7* cKO mice (Figures 1D, 1E). These findings suggest that the loss of Prmt7 preferentially affects CD8⁺ T cells over CD4⁺ T cells in the spleen. Consistently, our RT-qPCR data show that CD8⁺ T cells express ∼4-fold higher *Prmt7* mRNA levels compared to CD4⁺ T (Figure 1F and Supplemental Figure S1C).

**Figure 1:**
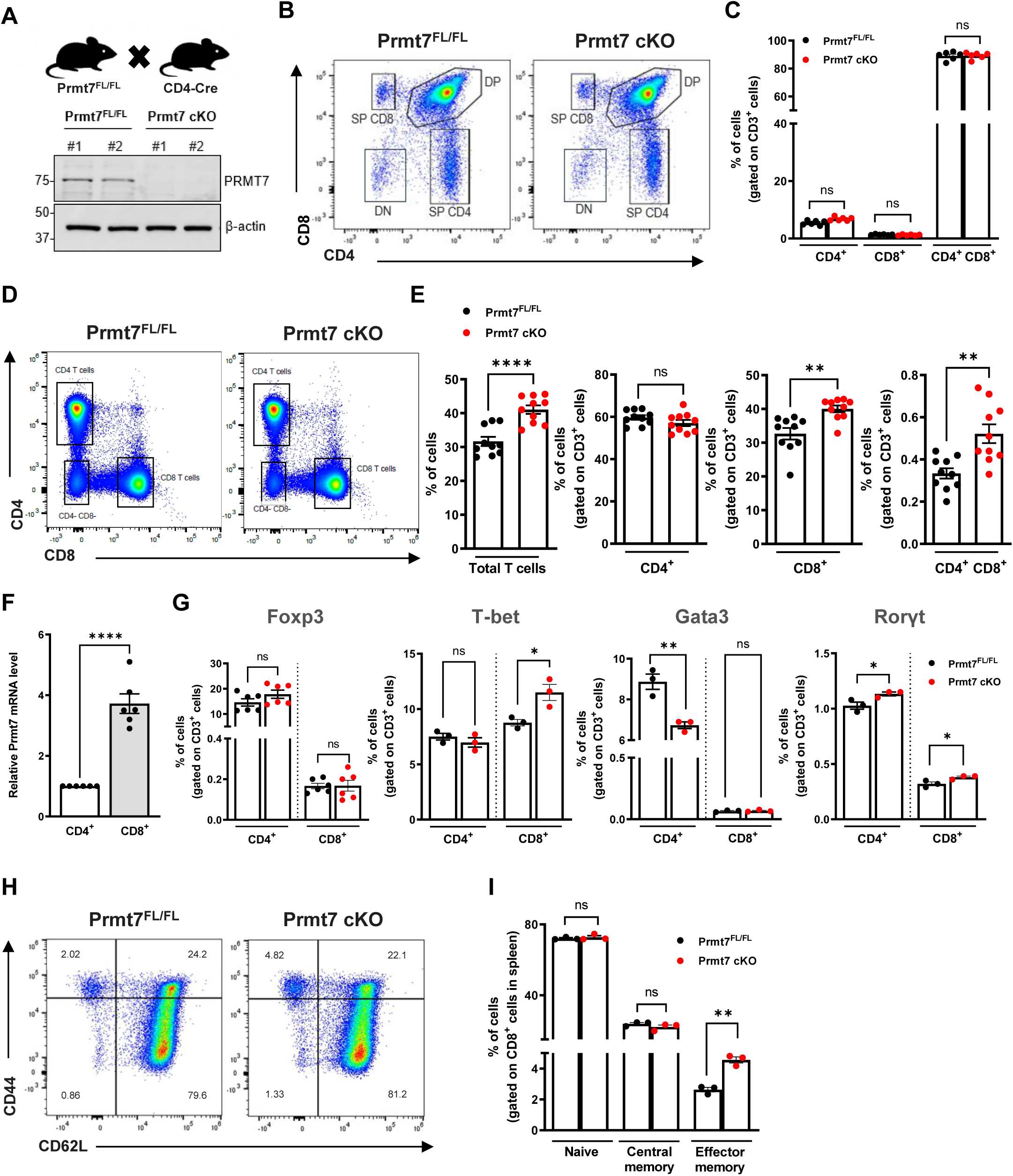
*Prmt7^FL/FL; CD4-Cre^* mice have increased CD8^+^ T cells in the periphery. **A.** Western blot showing Prmt7 expression in thymocytes from *Prmt7^FL/FL; CD4-Cre^* (*Prmt7* cKO) and littermate control *Prmt7^FL/FL^* mice. β-actin was used as the loading control. Molecular mass markers are indicated on the left in kDa. Data from two independent experiments are shown, each corresponding to a different mouse (#1, #2). **B**. Representative flow cytometry plots showing CD4 and CD8 expression in thymocytes isolated from *Prmt7^FL/FL^* control and *Prmt7* cKO mice. Cells were gated as indicated. **C.** Quantification graphs from **(B)** showing the percentages of single-positive SP (CD4^+^) and (CD8^+^), and double positive DP (CD4^+^ CD8^+^) thymic T cell subsets (gated on CD3^+^ cells) in *Prmt7^FL/FL^* control (black dots) and *Prmt7* cKO mice (red dots) (n=6). **D.** Representative flow cytometry plots showing CD4 and CD8 expression in splenocytes isolated from *Prmt7^FL/FL^* control and *Prmt7* cKO mice. Cells were gated as indicated. **E.** Quantification graphs from **(D)** showing the percentages of total splenic T cells, SP CD4^+^ and CD8^+^, and DP (CD4^+^ CD8^+^) T cell subsets (gated on CD3^+^ cells) in *Prmt7^FL/FL^* control (black dots) and *Prmt7* cKO mice (red dots) (n=10). **F.** Relative *Prmt7* mRNA levels quantified by RT-qPCR in CD4^+^ (empty bar) and CD8^+^ (grey bar) T cells sorted from wild-type mouse spleens (n=6). **G.** Percentages of FoxP3⁺, T-bet⁺, Gata3⁺ and Rroγt⁺ cells within gated CD4⁺ and CD8⁺ populations in *Prmt7^FL/FL^* control (black dots) and *Prmt7* cKO mice (red dots). (n=3-6). **H.** Representative flow cytometry plots showing CD62L and CD44 expression in splenocytes isolated from control and *Prmt7* cKO mice within gated CD8⁺ populations **I.** Percentage of CD62L^+^CD44^-^ (naïve), CD62L^+^CD44^+^ (central memory) and CD62L^-^CD44^+^ (effector memory) T cell subsets gated on CD8^+^ T cells, in *Prmt7^FL/FL^* control (black dots) and *Prmt7* cKO (red dots) mice (n=3). Data are representative of two to three independent experiments. Means ± SEM are shown. Statistical significance was determined using unpaired Student’s *t*-test (\**p* <0.05; \*\**p* <0.01; \*\*\**p* <0.001; \*\*\*\**p* <0.0001; *ns*: not significant).

Analysis of T cell subsets in the spleen revealed no significant changes in FoxP3 expression, suggesting that Prmt7 deficiency does not alter the differentiation of regulatory T cells (Tregs) (Fontenot *et al*, 2003) (Figure 1G). However, Prmt7-deficient mice showed an increase in CD8⁺ T cells expressing T-bet and a decrease in CD4⁺ T cells expressing Gata3, suggesting a shift toward a Th1-dominant immune profile (Szabo *et al*, 2000; Zheng & Flavell, 1997). In addition, expression of Rorγt was elevated in both CD4^+^ and CD8^+^ T cells (Figure 1G), suggesting enhanced Th17/Tc17 polarization (Ivanov *et al*, 2006).

Next, we profiled CD4^+^ and CD8^+^ T cells for naïve, central or effector memory, by staining for CD62L and CD44 markers. We observed a significant increase in the number of effector memory CD8^+^ cells harboring CD62L^lo^, CD44^hi^ in *Prmt7* cKO mice (Figure 1H, 1I), while no such increase was observed with the CD4^+^ T cells (Supplemental Figures S1D, S1E). These findings suggest that the loss of Prmt7 had a greater effect on CD8^+^ than CD4^+^ T cell populations, expanding their effector cell properties. In addition, *Prmt7* cKO mice exhibited higher frequencies of natural killer (NK) cells in the spleen (Supplemental Figures S1F, S1G).

### Prmt7 deficiency increases CD8^+^ T effector cell cytokine production and cytolytic activity

CD3^+^ T cells were isolated from spleens of *Prmt7^FL/FL^* (control) and *Prmt7* cKO mice to investigate their proliferation following activation with αCD3/CD28 beads. The proliferation index was assessed by flow cytometry after 3 days in culture through carboxyfluorescein succinimidyl ester (CFSE) dilution. Our data showed that *Prmt7*-deleted T cells exhibited ∼15% higher proliferation than control T cells (Figures 2A, 2B). After 3 days of activation, the CD8^+^ T cell population expanded markedly from 44.7% in *Prmt7^FL/FL^* to 67.1% in *Prmt7* cKO, while the CD4⁺ T cell population showed only a modest decrease (Figure 2C).

**Figure 2:**
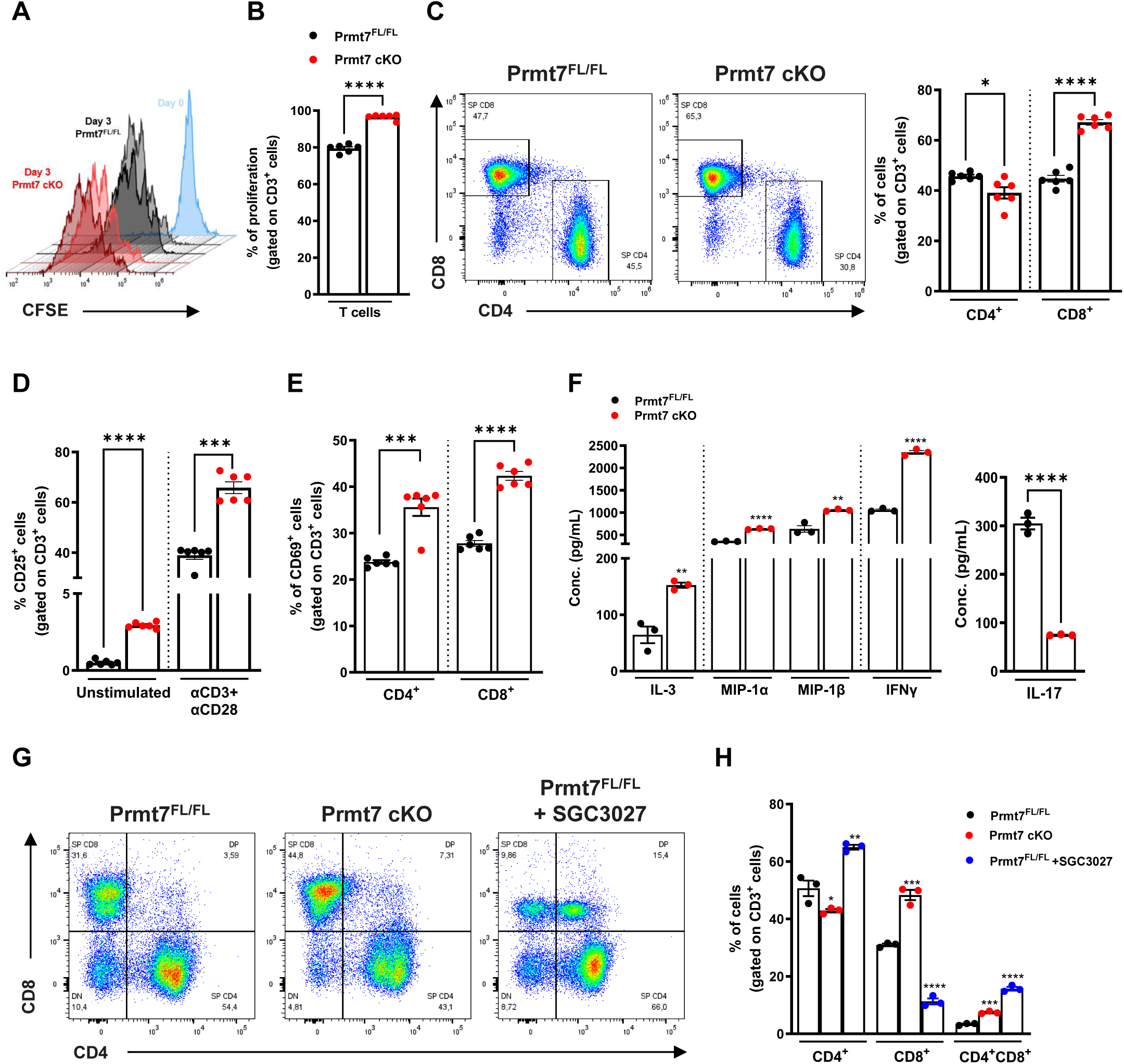
Prmt7 deficiency increases CD8^+^ T effector cell cytokine production and cytolytic activity. **A.** Freshly isolated T cells from *Prmt7^FL/FL^* control and *Prmt7* cKO (n=6) mice were obtained using Pan-T cell isolation kit, labeled with CFSE (5 µM), and stimulated with αCD3/CD28 Abs for 72h. The histograms represent unstimulated cells (blue, D0), stimulated cells from *Prmt7^FL/FL^* control mice (black, D3) and stimulated cells from *Prmt7* cKO mice (red, D3). One representative experiment is shown. **B.** Proliferation index was assessed by flow cytometry after 3 days in culture, based on CFSE dilution. The percentage of proliferating cells gated on CD3^+^ T cells is presented for *Prmt7^FL/FL^*control (black dots) and *Prmt7* cKO (red dots) mice (n=6). **C.** Representative flow cytometry plots showing CD4 and CD8 expression in splenocytes stimulated with αCD3/CD28 Abs for 72h. T cells were isolated from *Prmt7^FL/FL^*control (black dots) and *Prmt7* cKO (red dots) mice (n=6). The percentages of CD4^+^ and CD8^+^ T cell subsets, gated on CD3^+^ T cells, are shown. **D.** The percentage of T-cell activation using CD25 expression in unstimulated and stimulated T cells from *Prmt7^FL/FL^* control (black dots) and *Prmt7* cKO (red dots) mice (n=6). **E.** The percentage of T-cell activation using CD69 expression, gated on CD4^+^ and CD8^+^ T cells 72h post-stimulation from *Prmt7^FL/FL^*control (black dots) and *Prmt7* cKO mice (red dots) (n=6). **F.** Cytokine and Chemokine 44-plex assay measuring concentrations (pg/mL) of IL-3, MIP-1α, MIP-1β,IFN-γ and IL-17 in culture supernatants from *Prmt7^FL/FL^* control (black dots) and *Prmt7* cKO (red dots) T cells stimulated with anti-CD3/CD28 for 72h. **G.** Representative flow cytometry plots showing CD4 and CD8 expression in splenocytes stimulated with αCD3/CD28 Abs for 72h. T cells were isolated from *Prmt7^FL/FL^*control, *Prmt7* cKO mice and *Prmt7^FL/FL^* control mice treated with the PRMT7 inhibitor SGC3027 (10 μM). **H.** Quantification graphs from (**G)** showing the percentage of SP CD4^+^, SP CD8^+^ and DP CD4^+^CD8^+^ T cell subsets, gated on total CD3^+^ T cells, in the stimulated splenocytes from *Prmt7^FL/FL^*control (black dots), *Prmt7* cKO (red dots) mice and SGC-3027-treated splenocytes (blue dots) (n=3). Data are representative of 2-3 independent experiments. Means ± SEM are shown. Statistical significance was determined using unpaired Student’s *t*-test (\**p* <0.05; \*\**p* <0.01; \*\*\**p* <0.001; \*\*\*\**p* <0.0001; *ns*: not significant).

Consistently, flow cytometry revealed increased expression of the activation markers CD25 and CD69 in *Prmt7* cKO T cells under both unstimulated and stimulated conditions (Figures 2D, 2E), indicating a reprogrammed effector phenotype in the absence of Prmt7. This enhanced activation was accompanied by increased cytokine production, including IFNγ, IL-3, MIP-1α, and MIP-1β, in the culture supernatant of T cells from *Prmt7* cKO mice, whereas IL-17 levels were reduced, suggesting that Prmt7 deficiency promotes functional CD8⁺ T cell responses (Figure 2F).

To further determine whether the PRMT7 enzymatic activity is required for the T cell expansion observed, splenic CD3^+^ T cells isolated from control *Prmt7^FL/FL^* mice were treated with the PRMT7 inhibitor, SGC3027 (Szewczyk *et al*., 2020). Unlike CD8^+^ cells from *Prmt7* cKO, which showed an increase from ∼30% to ∼48% (Figures 2G, 2H, bars with red dots), SGC3027 treatment did not promote CD8⁺ expansion. Instead, a drastic decrease was observed in SGC3027-treated CD8^+^ T cells, dropping from ∼30% to ∼11% (Figures 2G, 2H, bars with blue dots), accompanied by a modest increase in CD4⁺ T cells. These results indicate that the increase in CD8^+^ T cells observed in *Prmt7* cKO mice is likely due to a non-catalytic function of PRMT7.

### Prmt7-deficiency leads to CD8^+^ T cell transcriptional reprogramming

To understand the molecular mechanism by which Prmt7 regulates CD8^+^ T cell development in the periphery, we analyzed the RNA profiles of purified CD8^+^ T cells from *Prmt7^FL/FL^* and *Prmt7* cKO mice (Figure 3 and Supplemental Figure S2). Gene Ontology (GO) analysis of the CD8^+^ T cell differential gene expression profiles (upregulation and downregulation; FDR <0.05) are consistent with activated cytotoxic T cells (CTLs) with enrichment in T cell receptor regulation, interleukin-2 signaling pathway, and T cell-specific transcription factors (TFs), and a downregulation of programmed cell death and apoptosis processes in *Prmt7* cKO mice (Figure 3A and Supplemental Figures S2A, S2B). A close examination of the major TF pathways activated in PRMT7-deficient T cells revealed increased activity of STAT5A, RELA, STAT3, MYC, ATF4, ATF5, CREB, FOS, ELK and SRF (Figure 3B and Supplemental Figure S2C), suggesting transcriptional reprogramming. These TFs are important because they are known to promote T cell activation, survival, proliferation, and cytokine production. Interestingly, we also observed a significant decrease in NAB2 repressor activity in Prmt*7* cKO T cells (Figure 3B). Reduced NAB2 activity may alleviate transcriptional repression of genes that drive T cell activation and effector function, suggesting that Prmt7 deletion favors a more activated and responsive T cell state. In addition, our analysis revealed that RelA ranked 9^th^ (p-value =5.56e^-17^) among the most variable TFs, showing a strong concordance with changes in TF activity between T cells from *Prmt7^FL/FL^* and *Prmt7* cKO mice (Figure 3B and Supplemental Figures S2D, S2E). Furthermore, a heat map plot showed that *NF-κB* gene expression levels (*NF-κB1, NF-κB2*, *RelA*, *RelB*) were higher in T cells isolated from *Prmt7* cKO compared to *Prmt7^FL/FL^*mice and this was confirmed by RT-qPCR (Figures 3C, 3D). Moreover, we validated the increased expression of several NF-κB target genes including iNos, ICAM, Bcl-2, Bcl-xL, and CD25 in CD8^+^ T cells using RT-qPCR (Figure 3D).

**Figure 3.**
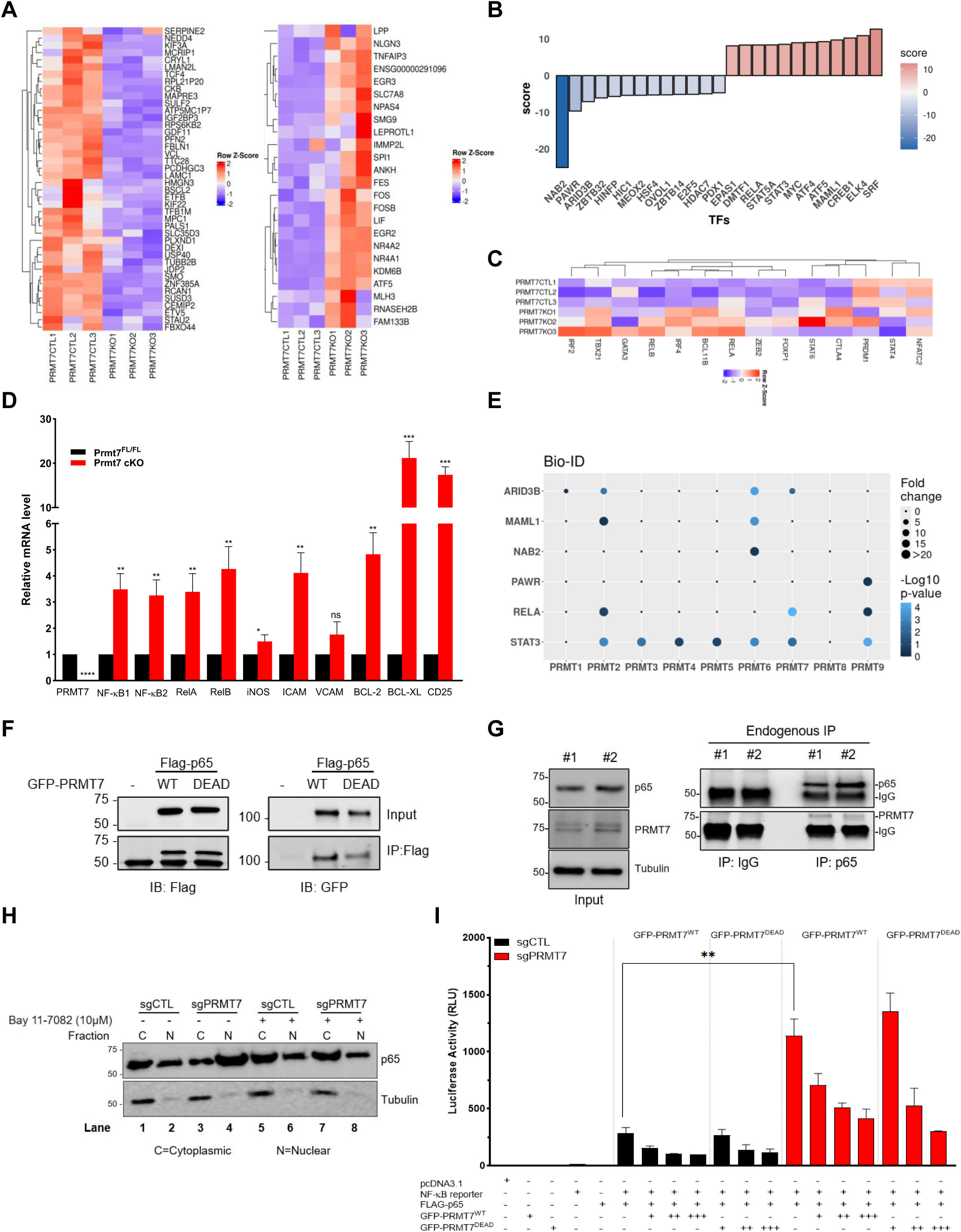
Prmt7-deficiency leads to CD8^+^ T cell transcriptional reprogramming. **A.** CD8^+^ T cells were isolated from mouse spleen and subjected to RNA-seq. Heatmaps depicting overlapping expression values (z-score based on cufflink count) of the top down and up-regulated genes between control (PRMT7CTL) and *Prmt7* cKO T cells (PRMT7KO). **B.** Changes in transcription factor (TF) activity, highlighting TFs with the largest differences in activity score between PRMT7CTL and PRMT7KO. Red bars indicate TFs that are activated in PRMT7KO compared to PRMT7CTL, while blue bars represent repressed TFs. **C.** Heatmaps showing overlapping expression patterns of NF-κB target genes between control and *Prmt7* cKO T cells. **D.** RT-qPCR analysis of direct and indirect NF-κB target gene expression including NF-κB1, NF-κB2, RelA, RelB, iNos, Icam, Vcam, Bcl2, Bcl-xL and CD25, in CD8^+^ T cells isolated from *Prmt7^FL/FL^* control mice (black bars) and *Prmt7* cKO mice (red bars). Data are representative of three independent experiments. Means ± SEM are shown. Statistical significance was determined using unpaired Student’s *t*-test (\**p* <0.05; \*\**p* <0.01; \*\*\**p* <0.001; \*\*\*\**p* <0.0001; *ns*: not significant). **E.** BioID analysis of PRMTs interactors. Dot size represents fold change in enrichment, while color intensity reflects statistical significance (-Log10 p-value), highlighting PRMTs-specific interaction patterns. **F.** B16.F10 cells were co-transfected with expression plasmids encoding Flag-p65 and GFP-PRMT7 (wild type, PRMT7^WT^ or catalytically inactive mutant, PRMT7^DEAD^). The cells were lysed and immunoprecipitated with anti-Flag beads. The bound proteins were separated by SDS-PAGE and subsequently immunoblotted with anti-GFP and anti-Flag antibodies. **G.** Mouse T cells were lysed and p65 immunoprecipitations performed. The presence of co-immunoprecipitating PRMT7 was detected by immunoblotting. **H.** sgCTL and Prmt7 Crispr-KO B16.F10 melanoma cells (sgPrmt7) were treated with the NF-κB inhibitor BAY 11-7082 (10μM) for 6h. Following treatment, Cytoplasmic (C) and Nuclear (N) fractions were prepared from treated (+) and untreated (-) cells and analyzed by immunoblotting using an anti-p65 antibody. Tubulin was used as a loading control for the cytoplasmic fractions. Molecular mass markers (kDa) are indicated on the left. Data are representative of two independent experiments. **I.** sgCTL and sgPrmt7 B16.F10 melanoma cells were transfected with pRLTK control plasmid (100 ng), pGL3 promoter NF-κB luciferase reporter plasmid (200 ng) and FLAG tagged p65 expression plasmid (400 ng) together with an increase amount of GFP tagged PRMT7^WT^ or the catalytically inactive mutant GFP-PRMT7^DEAD^ expression plasmids: 0 ng (-), 80 ng (+), 400 ng (++) and 2000 ng (+++). In all transfections, the pcDNA3.1 vector was added to bring the total plasmid to the same amount. Luciferase activity was analyzed 24h post-transfection using the Dual-Luciferase Reporter assay (Promega). Relative luciferase activity (RLU) was measured relative to the basal level of reporter gene in the presence of pcDNA3.1 vector after normalization with co-transfected RLU activity. Values mean ± SD for three independent experiments. Statistical significance was determined using unpaired Student’s *t*-test (\*\**p* <0.01).

Since the loss of PRMT7, rather than just inhibition of its activity by SGC3027, has significant effects on T cell function, we next investigated PRMT7 potential protein interactors. To do this, we analyzed previously generated PRMT7 BioID/mass spectrometry (MS) data (Wei *et al*, 2021) to identify relevant interactions that might modulate TF pathways. Surprisingly, only six TFs overlapped between the BioID dataset, and the top TFs identified by decoupleR from our RNA-seq analysis were ARID3B, MAML1, NAB2, PAWR, RELA, and STAT3. Our analysis revealed that among these six TFs, only RelA showed a high-confidence interaction with PRMT7 based on average intensity across three replicates, fold-change from mock control, and *p* values comparing mock control with the corresponding PRMT (Wei *et al*., 2021) (Figure 3E). AlphaFold modeling predicts an interaction (iPTM =0.22) between p65 and PRMT7, where 2 beta sheets of N-terminus of PRMT7 (in yellow) stack with the beta sheets of the dimerization domain of RelA (cyan) (Supplemental Figure S2F).

To confirm the interaction between PRMT7 and RelA also called p65, B16.F10 melanoma cells were transiently transfected with expression vectors encoding FLAG-epitope tagged p65 (FLAG-p65) and green fluorescent protein (GFP)- tagged PRMT7^WT^ or PRMT7^DEAD^, enzymatically inactive. Cell lysates were subjected to anti-FLAG immunoprecipitations, and bound proteins were analyzed by immunoblotting with anti-GFP antibodies. Both GFP-PRMT7^WT^ and GFP-PRMT7 ^DEAD^ co-immunoprecipitated equally with FLAG-p65 when adjusted to input (Figure 3F). These findings suggest that the methyltransferase activity of PRMT7 was not necessary for interaction with p65. Next, we proceeded with immunoprecipitation of endogenous proteins and indeed Prmt7 bound p65 in T cells, as Prmt7 was found in anti-p65 antibody immunoprecipitations, but not in the control IgG immunoprecipitations, performed in duplicate (Figure 3G). Collectively, these findings show that Prmt7 associates with p65 in T cells.

To understand how Prmt7 regulates the NF-κB pathway, we first explored the possibility that PRMT7 loss facilitates the nuclear translocation of p65. For this purpose, we took advantage of our Prmt7 Crispr-KO B16.F10 melanoma cells (sgPRMT7) (Srour *et al*., 2022), and treated them with the NF-κB inhibitor BAY 11-7082 at 10μM for 6h. Subsequently, cells were fractionated into nuclear (N) and cytoplasmic (C) fractions, followed by western blot analysis with an anti-p65 antibody. Indeed, we observed an increase in nuclear p65 in sgPRMT7 cells (Figure 3H, lane 4), which was reversed with BAY 11-7082 treatment (Figure 3H, lanes 6 and 8). Tubulin served as a control to confirm the absence of cytoplasm contamination in the nuclear fraction (Figure 3H). In addition, we performed immunofluorescence staining in sgCTL and sgPRMT7 B16.F10 melanoma cells following TNFα treatment known to activate the NF-κB pathway and Prmt7-deficient cells exhibited enhanced p65 nuclear staining compared to control cells (Supplemental Figure S3A), consistent with increased NF-κB activation.

Next, we overexpressed GFP-PRMT7 in wild type B16.F10 melanoma cells and examined p65 localization by indirect immunofluorescence. Overexpression of PRMT7 (green cells; GFP^+^) showed higher p65 cytoplasmic staining compared to non-transfected cells (GFP^-^) (Supplemental Figure S3B) as quantified in Supplemental Figure S3C, suggesting that PRMT7 retained p65 (RelA) in the cytoplasm. We noted that GFP-PRMT7 was predominantly localized in the cytoplasm of the B16.F10 melanoma cells (Supplemental Figure S3B) and we show by cell fractionation that Prmt7 was mainly cytoplasmic in T cells (Supplemental Figure S3D).

To directly assess whether Prmt7 influences NF-κB-mediated transcription, we performed a luciferase reporter assay in B16.F10 melanoma cells expressing p65. PRMT7-deficient cells (red bars) exhibited a ∼3-fold increase in NF-κB luciferase activity compared to control cells (black bars), suggesting that Prmt7 represses p65 transactivation (Figure 3I). Increasing amounts of expression vectors encoding GFP-PRMT7^WT^ or GFP-PRMT7^DEAD^ into sgCTL and sgPRMT7 B16.F10 melanoma cells repressed the p65-mediated luciferase activity, in a dose-dependent manner, suggesting that Prmt7 represses NF-κB activity in a methyltransferase-independent manner (Figure 3I).

### PRMT7 modulates the CD8^+^ T cell response and anti-tumor activity *via* the NF-κB pathway

We then investigated whether the observed increased proliferation and activation of CD8^+^ T cells was dependent on the NF-κB pathway. To examine this, we stimulated T cells from *Prmt7^FL/FL^* and *Prmt7* cKO mice with αCD3/CD28 activation beads for 3 days and treated them with the NF-κB inhibitor BAY 11-7082 for the last 6 hours. Flow cytometry analysis revealed that the increased CD8^+^ T cell population in *Prmt7* cKO mice was abolished with BAY 11-7082 (Figures 4A, 4B). These results indicate that the enhanced percentage of CD8^+^ T cells in *Prmt7* cKO mice was dependent on NF-κB signaling.

**Figure 4.**
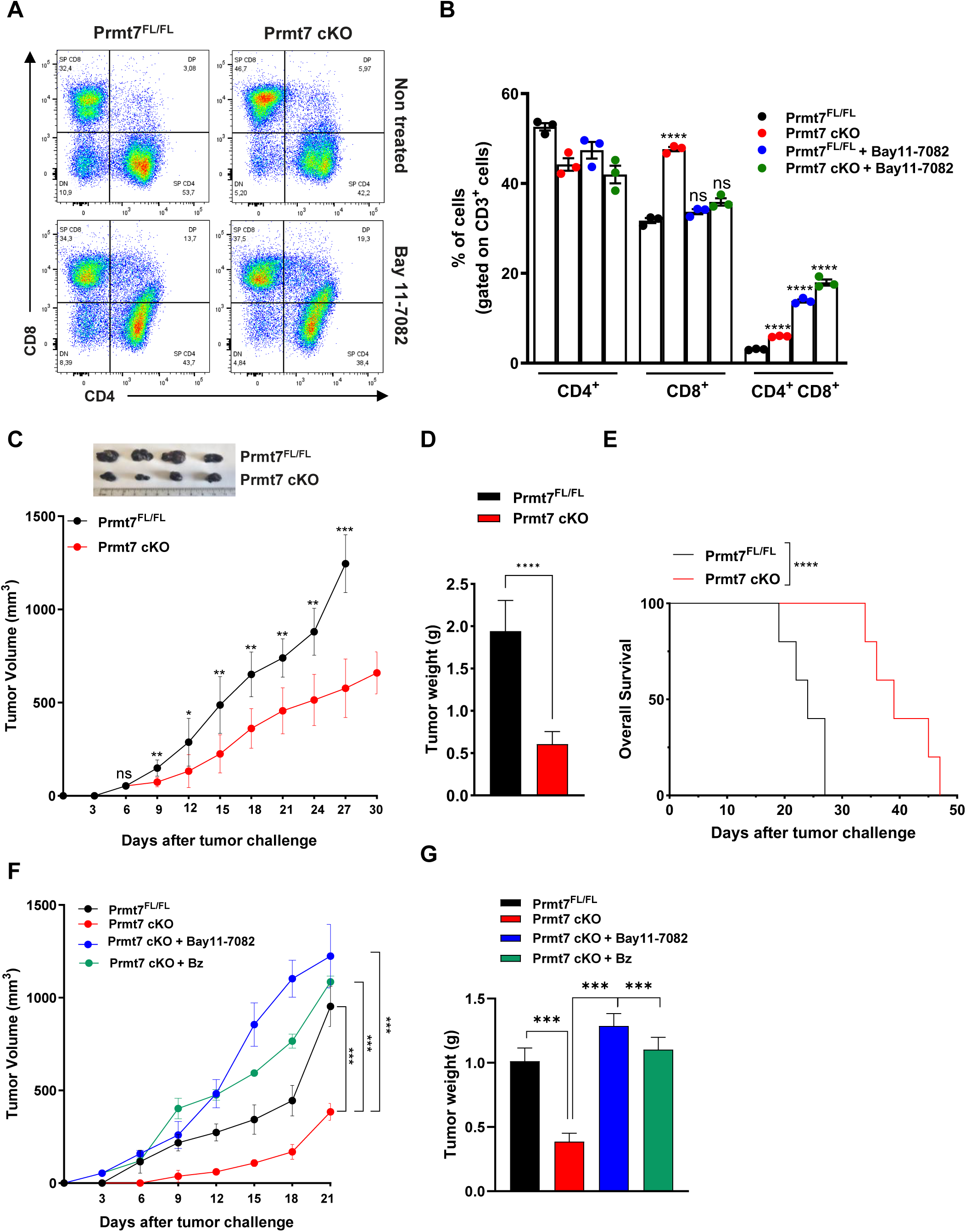
PRMT7 modulates CD8^+^ T cell responses and anti-tumor activity *via* the NF- κB pathway. **A.** Representative flow cytometry plots showing CD4 and CD8 expression in splenocytes from *Prmt7^FL/FL^* control and *Prmt7* cKO mice. Splenocytes were stimulated with αCD3/CD28 Abs for 72h followed by treatment with the NF-κB inhibitor BAY 11-7082 (10μM) for the last 6h. **B.** Quantification graphs from panel **(A)**, showing the percentages of SP CD4^+^, SP CD8^+^ and DP (CD4^+^ CD8^+^) T cell subsets, gated on total CD3^+^ T cells. Data are presented for stimulated splenocytes from *Prmt7^FL/FL^* control (black dots) and *Prmt7* cKO (red dots) mice, as well as splenocytes treated with BAY 11-7082 (10μM) (blue and green dots) (n=3 mice/group). **C.** B16.F10 melanoma cells were injected subcutaneously into *Prmt7^FL/FL^*control and *Prmt7* cKO mice, and representative images of tumors are shown. Tumor volumes were averaged for each group at each time point (*Prmt7^FL/FL^*, black; *Prmt7* cKO, red). Tumor growth curves were analyzed using two-way ANOVA. (\**p* <0.05; \*\**p* <0.01; \*\*\**p* <0. 001; *ns*: not significant). **D.** Tumor weight (g) at endpoint as observed in **(C)**, is shown. Mean ± SEM is presented. Statistical significance was determined using unpaired Student’s *t*-test (\*\*\*\**p* <0.0001). **E.** Kaplan-Meier survival curves were assessed at the indicated time points for the mice in **(C)**. n = 5 mice/group; *p* values were determined using multiple *t*-test (\*\*\*\**p* <0.0001). **F.** B16.F10 melanoma cells were subcutaneously injected into *Prmt7^FL/FL^* control (black) and *Prmt7* cKO mice (red) followed by intraperitoneal injection of either BAY 11-7082 (5mg/kg; blue) or Bortezomib (1mg/kg; green) in *Prmt7* cKO mice. Tumor volumes were averaged for each group at each time point (n=5 mice/group). Tumor growth curves were analyzed using two-way ANOVA. (\*\*\**p* <0. 001). **G.** Tumor weight (g) at endpoint as observed in **(F)**, is shown. Mean ± SEM is presented. Statistical significance was determined using unpaired Student’s *t*-test (\*\*\**p* <0.001).

We next proceeded to a tumor challenge in *Prmt7^FL/FL^* and *Prmt7* cKO mice using the B16.F10 melanoma syngeneic murine model. Interestingly, B16.F10 melanomas exhibited slower outgrowth, reduced tumor burden and increased survival of the *Prmt7* cKO mice, compared to control (Figures 4C-4E). These data suggest that the CD8^+^ T cells from the *Prmt7* cKO mice have enhanced effector function and increased cytotoxicity towards the melanoma *in vivo*. To further assess whether the tumor phenotype observed in *Prmt7* cKO mice was mediated by NF-κB signaling, we treated these mice with two distinct NF-κB inhibitors: BAY 11-7082 (5 mg/kg; i.p., blue bars) and Bortezomib (1 mg/kg; i.p., green bars). Treatment with either inhibitor, compared to vehicle, significantly increased tumor volume (Figure 4F) and weight (Figure 4G). Tumor infiltrating lymphocytes (TILs) analysis revealed that tumors from *Prmt7* cKO mice treated with BAY11-7082 or Bortezomib were markedly less infiltrated, showing reduced lymphoid and myeloid immune cells (Supplemental Figures S4A, S4B). Both treatments decreased T cells and myeloid subsets, indicating a colder tumor microenvironment and reduced immune infiltration compared with untreated *Prmt7* cKO tumors.

### Discovery of MS54, a PRMT7 PROTAC that increases CD8^+^ T effector cell cytokine production and cytolytic activity

The PRMT7 inhibitor, SGC3027, did not mimic our findings with the *Prmt7* cKO phenotype (Figures 2G, 2H), suggesting that suppression of Prmt7 expression, rather than blockade of its methyltransferase activity, was necessary to drive the anti-tumor CD8^+^ T cell response observed. Therefore, we proceeded to generate a PRMT7 PROTAC degrader, as we have previously generated for PRMT5 and PRMT6 (Shen *et al*., 2020a; Shen *et al*., 2020b). To develop the PRMT7 PROTAC, we first chose SGC8158 (Szewczyk *et al*., 2020), a potent and selective PRMT7 inhibitor, as the warhead for PRMT7 binding (Figure 5A). Analysis of the co-crystal structure of SGC8158 in complex with PRMT7 (PDB ID: 6OGN) revealed that the 4-position of its 1,1’-biphenyl moiety is solvent-exposed, providing a suitable site for linker attachment without disrupting PRMT7 binding (Supplemental Figure S5A). Based on this result, we introduced a 2-oxyethylamine group at the 4-position to serve as a handle for subsequent linker installation. Next, we conjugated this intermediate (SGC8158 amine, Figure 5A) with VHL101 acid (Figure 5A), a well-known VHL E3 ligase ligand (Diehl & Ciulli, 2022), via various linkers (alkyl or polyethylene glycol (PEG)), yielding a series of putative PRMT7 PROTAC degraders (compounds 1-13) (Supplemental Figure S5B). These degraders were screened by western blot analysis in B16.F10 mouse melanoma cells at two concentrations (3 and 10 μM) (Supplemental Figure S5C). Degraders with relatively short alkyl linkers (compounds 1-4) had minimal effects on Prmt7 expression. Beginning from compound 5, moderate Prmt7 degradation was observed with approximately 40% reduction at 10 μM. Further increasing the linker length enhanced degradation efficiency, with compound 8 (MS54) showing the highest potency, achieving over 80% degradation at 10 μM. Notably, despite the poor cellular permeability of the Prmt7 binder SGC8158, MS54 effectively degraded Prmt7 in cells, underscoring a key advantage of PROTAC degraders over conventional small-molecule inhibitors (Dale *et al*, 2021). In contrast, degraders with PEG linkers (compound 9-13) failed to induce significant Prmt7 degradation.

**Figure 5.**
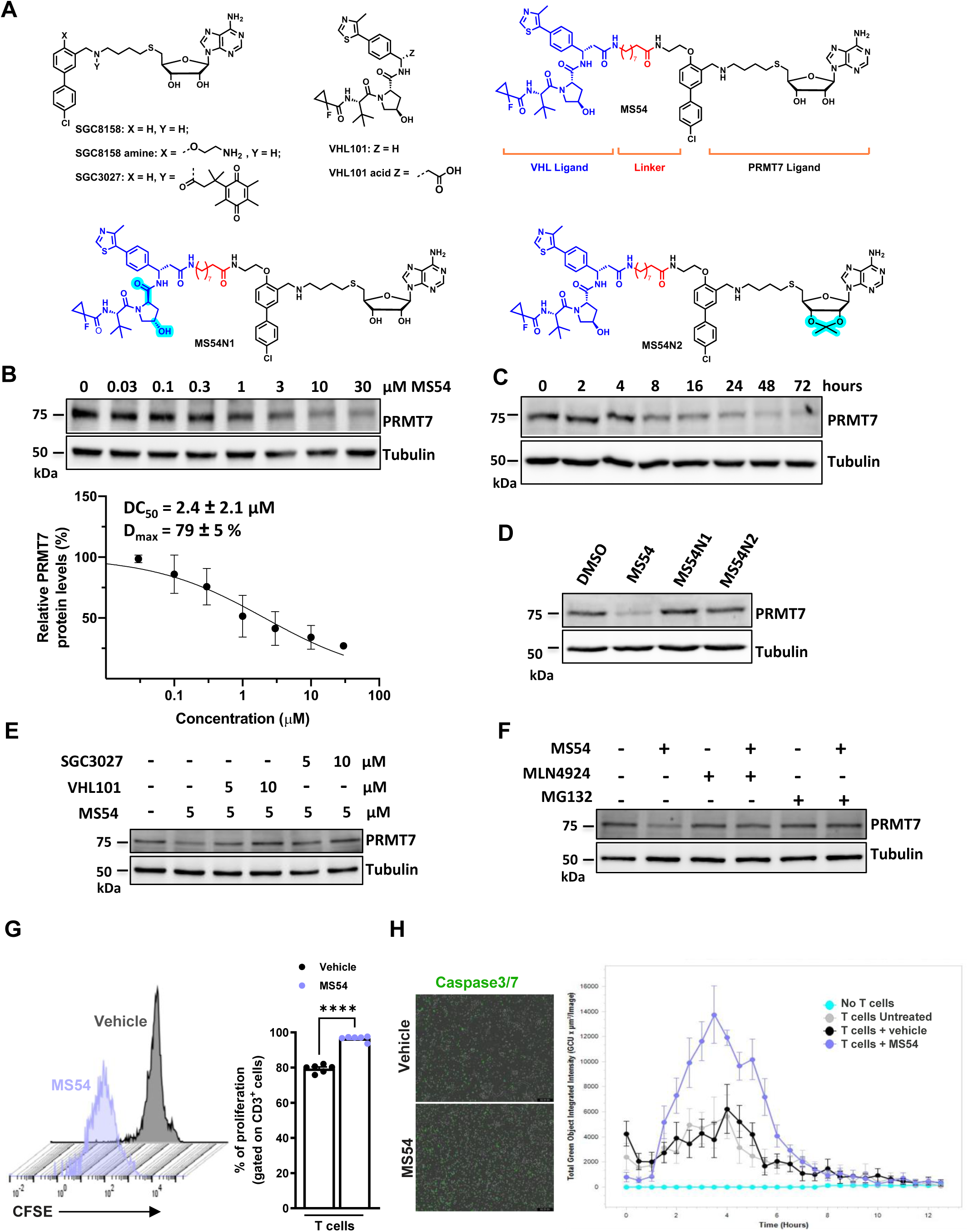
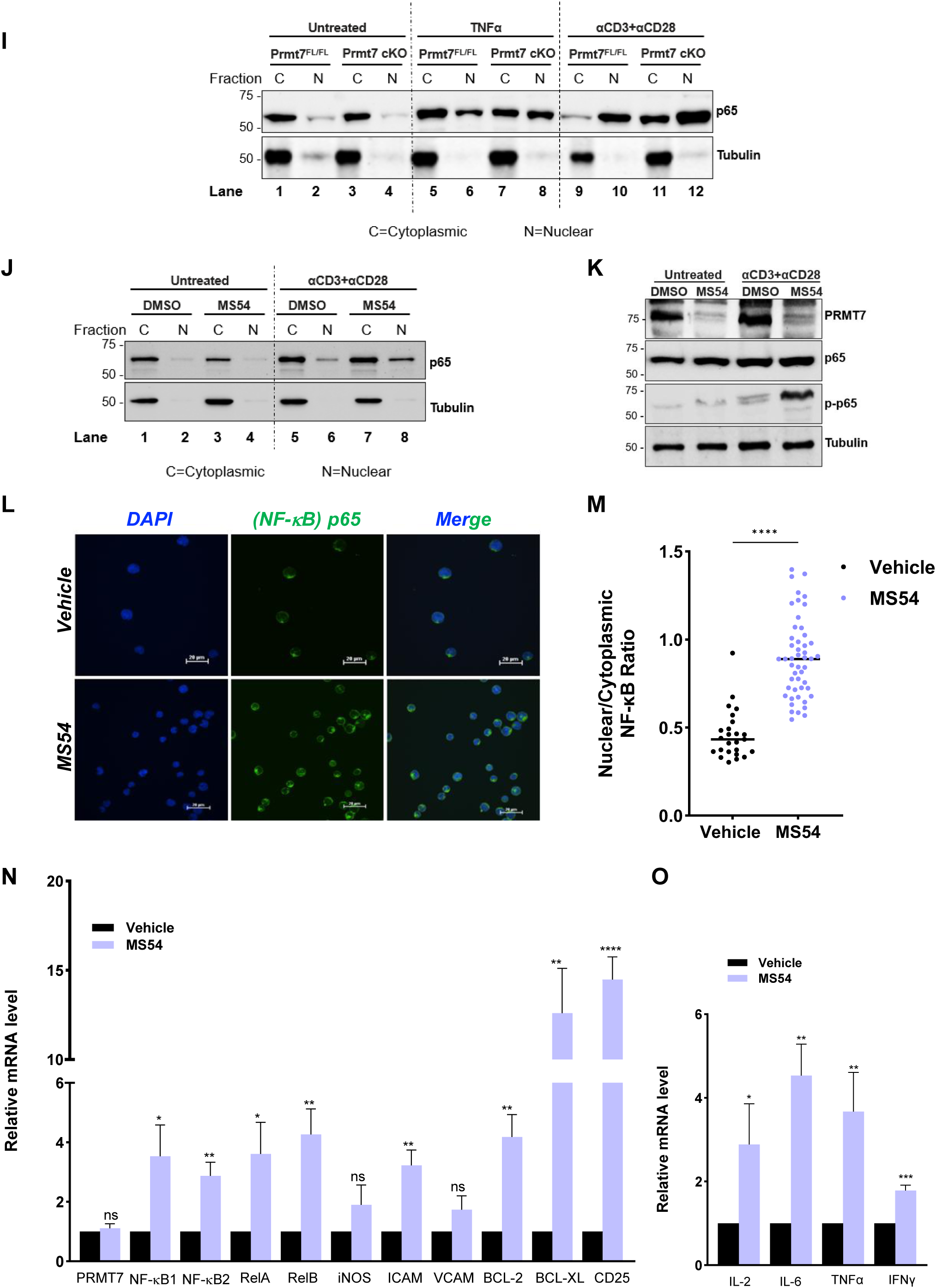
Discovery of MS54, a PRMT7 PROTAC that increases CD8^+^ T effector cell cytokine production and cytolytic activity. **A.** Chemical structures of SGC8158, SGC8158 amine, SGC3027, VHL101, VHL101 acid, MS54, MS54N1 and MS54N2. **B.** Top, immunoblot of PRMT7 in B16.F10 cells treated with the indicated concentration of MS54, versus DMSO, for 24 h. Bottom, measurement of DC_50_ and D_max_ values of MS54, based on the PRMT7 immunoblot signals after treatment. Mean ± SD is presented (n = 3). **C.** Time course of PRMT7 degradation by MS54 (10 μM) in B16.F10 cells. One representative image is shown. **D.** Immunoblot of PRMT7 in B16.F10 cells treated with the indicated compound (10 μM) for 24 h. One representative image is shown. **E.** Immunoblot of PRMT7 in B16.F10 cells pretreated with SGC3027, or VHL101 at indicated concentration for 2 h followed by MS54 (5 μM) for 24 h. One representative image is shown. **F.** Immunoblot of PRMT7 in B16.F10 cells pretreated with MG132 (10 μM), or MLN4924 (μM) for 2 h followed by MS54 (10 μM) for 24 h. One representative image is shown. **G.** Freshly isolated T cells from control (n=6) mice were treated with vehicle or 3 μM MS54. T cells were obtained using Pan-T cell isolation kit, labeled with CFSE (5 µM), and stimulated with αCD3/CD28 Abs for 72h. The histograms represent stimulated cells vehicle treated (black) or MS54 treated (purple). Proliferation index was assessed by flow cytometry after 3 days in culture, based on CFSE dilution. The percentage of proliferating cells gated on CD3^+^ T cells is presented for control (black dots) and *Prmt7* cKO (purple dots) mice (n=6). **H.** Real-time monitoring of cancer cell killing. Caspase-3/7 activation was measured using the IncuCyte® live-cell analysis system, with green fluorescence intensity (Caspase-3/7 dye; GCU) plotted on the Y-axis over time. B16.F10 cancer cells were co-cultured with pre-activated CD8⁺ T cells (48 h) at an effector-to-target (E:T) ratio of 10:1 under the following conditions: blue, negative control (no T cells); grey, T cells from *Prmt7^FL/FL^* (control mice); black, T cells treated with vehicle (DMSO); purple, T cells treated with MS54 (3 µM). Data are representative of three experiments and presented as means ± SEM. **I.** Western blot analysis of Cytoplasmic (C) and Nuclear (N) fractions showing increased nuclear accumulation of p65 in *Prmt7 cKO* T cells compared to *Prmt7^FL/FL^* controls following TNFα (6 h) or αCD3/CD28 (72 h) stimulation. **J.** Western blot analysis of Cytoplasmic (C) and Nuclear (N) fractions showing enhanced nuclear localization of p65 in T cells treated with MS54 (3 µM) and stimulated with αCD3/CD28 for 72 h. **(I-J).** Tubulin was used as a loading control for the cytoplasmic fractions. Molecular mass markers (kDa) are indicated on the left. Data are representative of two independent experiments. **K.** Immunoblot analysis of PRMT7, total p65 and phosphorylated p65 (p-p65) in T cells following 72 h αCD3/αCD28 stimulation with or without MS54 treatment (3 µM). Representative data from two independent experiments are shown. **L-M.** Representative immunofluorescence images **(L)** and quantification **(M)** of nuclear-to-cytoplasmic (N/C) fluorescence intensity ratios for p65 in DMSO- (black dots) and MS54-treated and activated human T cells (CCRF-CEM) (purple dots). Cells were imaged on a Nikon AXR point laser scanning confocal microscope and analyzed using GA3 (General Analysis 3) in NIS-Elements software. Magnification 60x and numerical aperture of the objective:1.42 NA. Means ± SEM are shown. Statistical significance was determined using unpaired Student’s *t*-test, \*\*\*\**p* <0.0001. **N.** RT-qPCR analysis of direct and indirect NF-κB target gene expression including NF-κB1, NF-κB2, RelA, RelB, iNos, Icam, Vcam, Bcl2, Bcl-xL and CD25, in T cells stimulated with αCD3/CD28 in the presence (purple bars) or absence (black bars) of MS54 (3µM). **O.** Relative mRNA expression levels of IL-2, IL-6, TNFα, and IFNγ in T cells stimulated with αCD3/CD28 in the presence (purple bars) or absence (black bars) of MS54 (3µM). Data are representative of three independent experiments. Means ± SEM are shown. Statistical significance was determined using unpaired Student’s *t*-test (\**p* <0.05; \*\**p* <0.01; \*\*\**p* <0.001; \*\*\*\**p* <0.0001; *ns*: not significant).

Following the identification of MS54 as the lead compound, it was further characterized through a series of biochemical and cellular assays. MS54 induced Prmt7 degradation in a concentration-dependent manner, with a DC50 of 2.4 ± 2.1 μM, and Dmax of 79 ± 5% (Figure 5B). Time-course studies showed that Prmt7 degradation began as early as 8 hours post-treatment, reached maximal degradation at 48 hours, and was sustained for up to 72 hours (Figure 5C). To elucidate the mechanism of action (MOA) of MS54, we designed two structurally related analogs, MS54N1 and MS54N2, as negative controls (Figure 5A). MS54N1 features an epimerized form of the VHL ligand VHL101, which disrupts VHL binding, while MS54N2 incorporates an isopropylidene-protecting group on the ribose moiety of SGC8158, intended to prevent PRMT7 binding without affecting VHL engagement. These compounds were further evaluated in the B16.F10 cell line. As shown in Figure 5D, only MS54, but not MS54N1 or MS54N2, induced Prmt7 degradation, indicating that the engagement of both PRMT7 and VHL is essential for MS54-mediated degradation. This was further supported by the rescue experiments. Pre-treatment with either SGC3027 (a prodrug of SGC8158) or VHL101 completely abolished MS54-induced Prmt7 degradation (Figure 5E). Moreover, pre-treatment with the proteasome inhibitor, MG132, (Lee & Goldberg, 1996) or the neddylation inhibitor, MLN4924, (Soucy *et al*, 2009) prevented the Prmt7 degradation (Figure 5F). Together with RT-qPCR analysis showing unchanged Prmt7 mRNA levels (Supplemental Figure S6A). These results strongly suggest that MS54 induced PRMT7 degradation *via* the ubiquitin-proteasome system (UPS).

To assess the impact of MS54 on T cell proliferation and activation, splenic T cells from *Prmt7^FL/FL^* mice were activated with αCD3/CD28 beads and treated with either vehicle (DMSO) or 3 µM MS54 for 72 hours. Proliferation index was assessed by flow cytometry after 3 days in culture through CFSE dilution (Figure 5G). Our data showed that MS54 treated T cells exhibited increased proliferation by ∼20% compared to untreated T cells (Figure 5G and Supplemental Figure S6B). After 3 days of activation, we observed a significant expansion of the CD8^+^ T cells from ∼15% in vehicle treated T cells to ∼30% in MS54 treated cells (Supplemental Figures S6C, S6D). Next, we assessed the activation markers CD25 and CD69 by flow cytometry in stimulated splenocytes and observed an increase in both markers in CD4⁺ and CD8⁺ T cell populations (Supplemental Figures S6E, S6F). These findings show that MS54 increases CD8⁺ T cell expansion, consistent with the phenotype observed in *Prmt7* cKO mice.

Next, we investigated whether MS54 treated CD8⁺ T cells exhibit enhanced cytotoxic activity. CD8⁺ T cells isolated from *Prmt7^FL/FL^* control mice were pre-activated and co-cultured with B16.F10 melanoma cells at an effector-to-target (E:T) ratio of 10:1. Tumor cell killing was assessed in real time using the IncuCyte® live-cell imaging system and Caspase-3/7 reagent (Sartorius, USA). MS54 treated CD8⁺ T cells displayed markedly increased cytotoxicity as evidenced by higher levels of apoptotic tumor cells (Figure 5H, left panel). Quantification of total green object integrated intensity (GCU) further confirmed elevated apoptosis in co-cultures with MS54-treated T cells (purple) compared to vehicle-treated controls (black) (Figure 5H, right panel). Together, these results demonstrate that MS54 enhances the cytotoxic capacity of CD8⁺ T cells against melanoma.

To assess whether MS54 influences NF-κB signaling during T cell activation, we examined the subcellular localization of the p65 subunit in control *Prmt7^FL/FL^* and *Prmt7* cKO T cells activated with TNFα (6 hours) or αCD3/CD28 (72 hours, Figure 5I). Western blot analysis of cytoplasmic and nuclear fractions revealed increased nuclear p65 in *Prmt7* cKO T cells compared to control following activation (Figure 5I). Similarly, treatment with MS54 resulted in Prmt7 degradation and increased p65 nuclear localization in T cells relative to DMSO-treated controls (Figure 5J). These data indicate that pharmacological degradation of Prmt7 by MS54 phenocopies the genetic knockout of Prmt7 in T cells. Moreover, western blot analysis of phosphorylated p65 (active form) revealed increased phosphorylation following T cell activation, which was further enhanced upon treatment with MS54 (Figure 5K).

Immunofluorescence staining further confirmed these findings, showing a significantly higher nuclear to cytoplasmic (N/C) fluorescence ratio of p65 in MS54-treated human CCRF-CEM T cells compared to their respective controls (Figures 5L, 5M). Upon activation, p65 nuclear translocation was markedly increased, consistent with enhanced NF-κB activation.

To determine whether this increased nuclear localization of p65 correlated with enhanced transcriptional output, we performed RT-qPCR analysis of NF-κB target genes in splenic T cells stimulated for 72 hours with αCD3/CD28 in the presence or absence of MS54. Our data show an increase in the expression of *NF-κB* genes and its target genes (Figure 5N). In addition, the expression of IL-2, IL-6, TNFα, and IFNγ, well-established NF-κB responsive cytokine genes, was significantly upregulated in MS54-treated T cells compared to controls (Figure 5O). Collectively, these results demonstrate that loss of PRMT7 promotes NF-κB activation in T cells, as evidenced by increased nuclear p65 translocation and elevated expression of downstream proinflammatory cytokine genes.

### Adoptive cell transfer (ACT) with MS54 treated T cells suppresses tumor growth

The enhanced generation of effector memory cytotoxic T lymphocytes (CTLs) and the increased expression of effector markers in CTLs led us to investigate whether MS54 pretreated CTLs *ex vivo* would have enhanced anti-tumor activity *in vivo* using ACT. To test this possibility, we isolated CD8^+^ T cells from OT-I transgenic mice, which express an MHC class I-restricted TCR specific for the OVA-derived SIINFEKL epitope. These OT-I T cells were activated *ex vivo* to become potent effector cells, in the presence of DMSO or 3 µM MS54 (Figure 6A). Activated OT-I T cells were adoptively transferred into C57BL/6J mice harbouring subcutaneous B16.F10-ovalbumin tumors.

**Figure 6.**
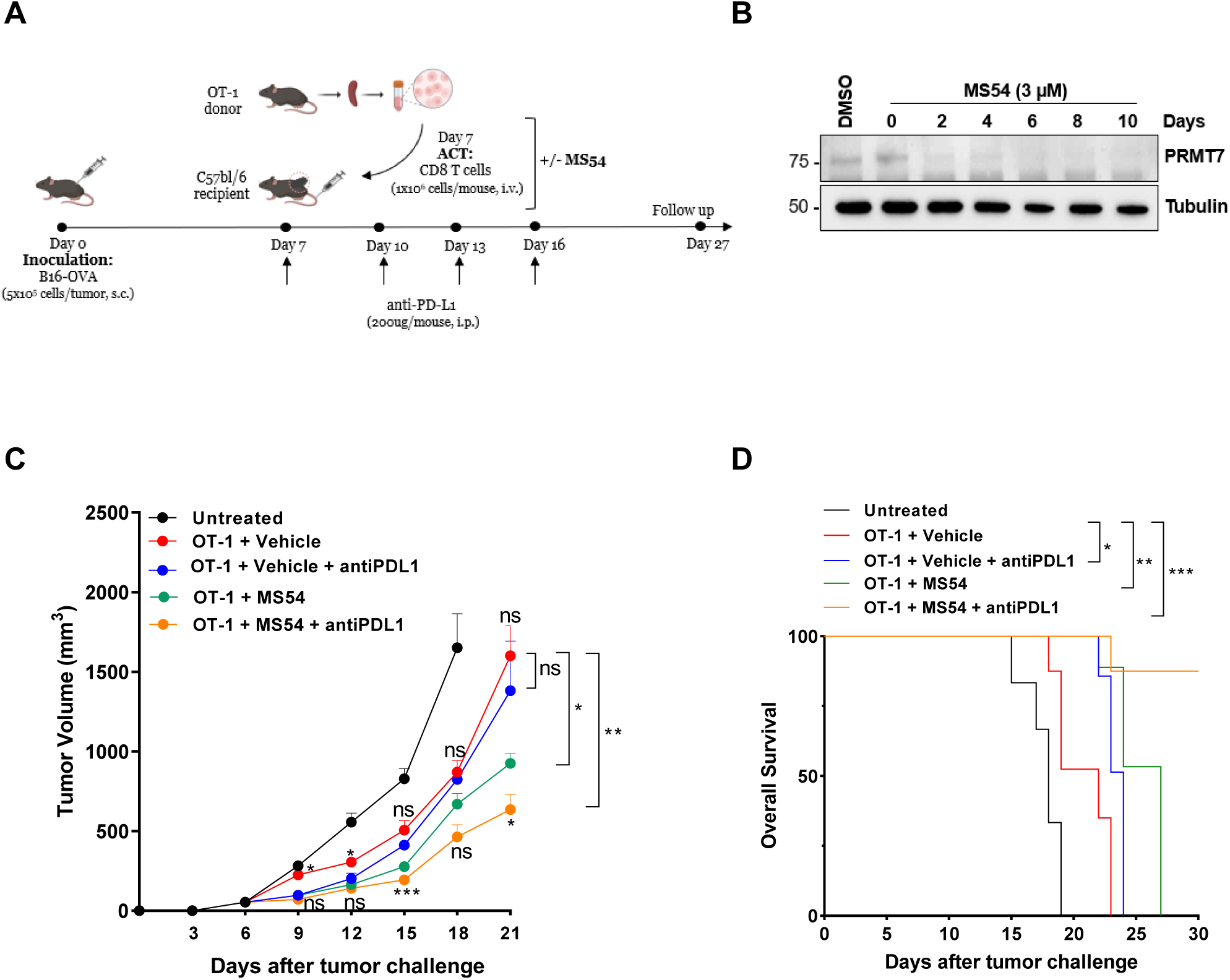
Adoptive cell transfer (ACT) with MS54-treated T cells suppresses tumor growth. **A.** Schematic of the experimental procedure. CD8^+^ T cells were isolated from OT-I mice, pre-activated *ex vivo* in the presence of IL-2, and treated with either DMSO or 3μM MS54 for 72h. On day 7, cells were then adoptively transferred into C57BL/6J mice (1×10^6^ cells/mouse i.v.), that were previously been implanted with B16.F10-OVA tumors cells (0.5×10^6^ cells/mouse s.c.). Mice then received either isotype control or anti-PD-L1 antibody (200 µg/mouse, i.p.) on days 7, 10, 13, and 16. **B.** Western blot showing PRMT7 degradation in CD8⁺ T cells treated with DMSO or 3 µM MS54 and activated *ex vivo* for up to 10 days. Tubulin served as a loading control. Molecular weight markers (kDa) are indicated on the left. A representative blot is shown. **C.** Tumor volume averaged for each group at each time point is presented. Data are mean ± SEM; n = 5 mice/group. Tumor growth curves were analyzed using two-way ANOVA (\**p* <0.05; \*\**p* <0.01; \*\*\**p* <0. 001; *ns*: not significant). **C.** Kaplan-Meier survival curves were assessed at the indicated time points for mice from panel **C**. n = 5 mice/group; *p* values were determined using multiple *t*-test (\**p* <0.05; \*\**p* <0.01; \*\*\**p* <0.001).

A separate cohort of mice received anti-PDL1 therapy (200 µg per mouse, equivalent to 10 mg/kg, i.p.) on days 7, 10, 13, and 16 following tumor implantations (Figure 6A). *Ex vivo* activated CD8⁺ T cells were treated with MS54 to assess the stability of Prmt7 degradation. MS54 efficiently reduced Prmt7 protein levels, and this effect persisted for over 10 days post-stimulation (Figure 6B).

Mice that received MS54 treated OT-I T cells exhibited markedly reduced tumor growth and improved survival compared to those receiving untreated OT-I effector cells (Figures 6C, 6D; red vs. green). Combination therapy with anti-PDL1 further enhanced survival in mice treated with MS54 (Figures 6C, 6D; blue vs orange). Notably, MS54-treated OT-I T cells alone conferred greater tumor control than untreated OT-I cells combined with anti-PDL1 therapy, underscoring the potency of Prmt7 degradation in enhancing CTL function. Together, these results demonstrate that the PRMT7 PROTAC degrader MS54 augments antigen-specific CD8⁺ T cell anti-tumor activity *in vivo*.

### MS54 enhances human CD8^+^ T lymphocyte effector functions

To evaluate the impact of PRMT7 degradation on human T cell function, we used an HLA-matched antigen recognition assay. Primary human CD8⁺ T cells engineered to express a transgenic NY-ESO-1-specific TCR (Gnjatic *et al*, 2006), were co-cultured with HLA-matched 624mel melanoma cells. Activation occurred in the presence of these cancer cells, which endogenously express NY-ESO-1, indicating an antigen-specific response (Figure 7A). These T cells were treated with DMSO or MS54 at 1 or 3μM for 72 hours and the expression of PRMT7 was assessed by immunoblotting (Figure 7B). In addition, we observed a significant increase in the number of MS-54-treated CD8⁺ T cells following coculture with 624mel cells, consistent with enhanced expansion (Figure 7C). Functionally, MS54-treated T cells exhibited increased expression of the activation markers CD69 and CD137 (Figure 7D and Supplemental Figure S7A), along with elevated IFN-γ secretion in the supernatants collected after 3 days of coculture with NY-ESO-1-specific T cells and 624mel cells, as measured by ELISA (Figure 7E). Representative FACS plots (Supplemental Figure S7B) and Mean Fluorescence Intensity (MFI) of Ki-67 labeling (Supplemental Figure S7C), demonstrated enhanced proliferative activity in NY-ESO-1-specific T cells co-cultured with 624mel cells when treated with 3 µM MS54 compared to untreated control. For functional killing assays, 624mel target cells were co-cultured with NY-ESO-1-specific T cells that had been pretreated with MS54 (1 or 3 µM) or DMSO for 24 h prior to coculture and the readout was performed 48 h later. Our data showed that MS54 pretreatment enhanced cytotoxic activity and tumor cell killing in a dose-dependent manner at various effector-to-target (E:T) ratios, with the strongest effect observed at an E:T ratio of 10:1 and a concentration of 3 µM MS54 (Figure 7F). Collectively, these findings demonstrate that PRMT7 degradation by MS54 enhanced antigen-specific effector responses in human CD8⁺ T cells.

**Figure 7.**
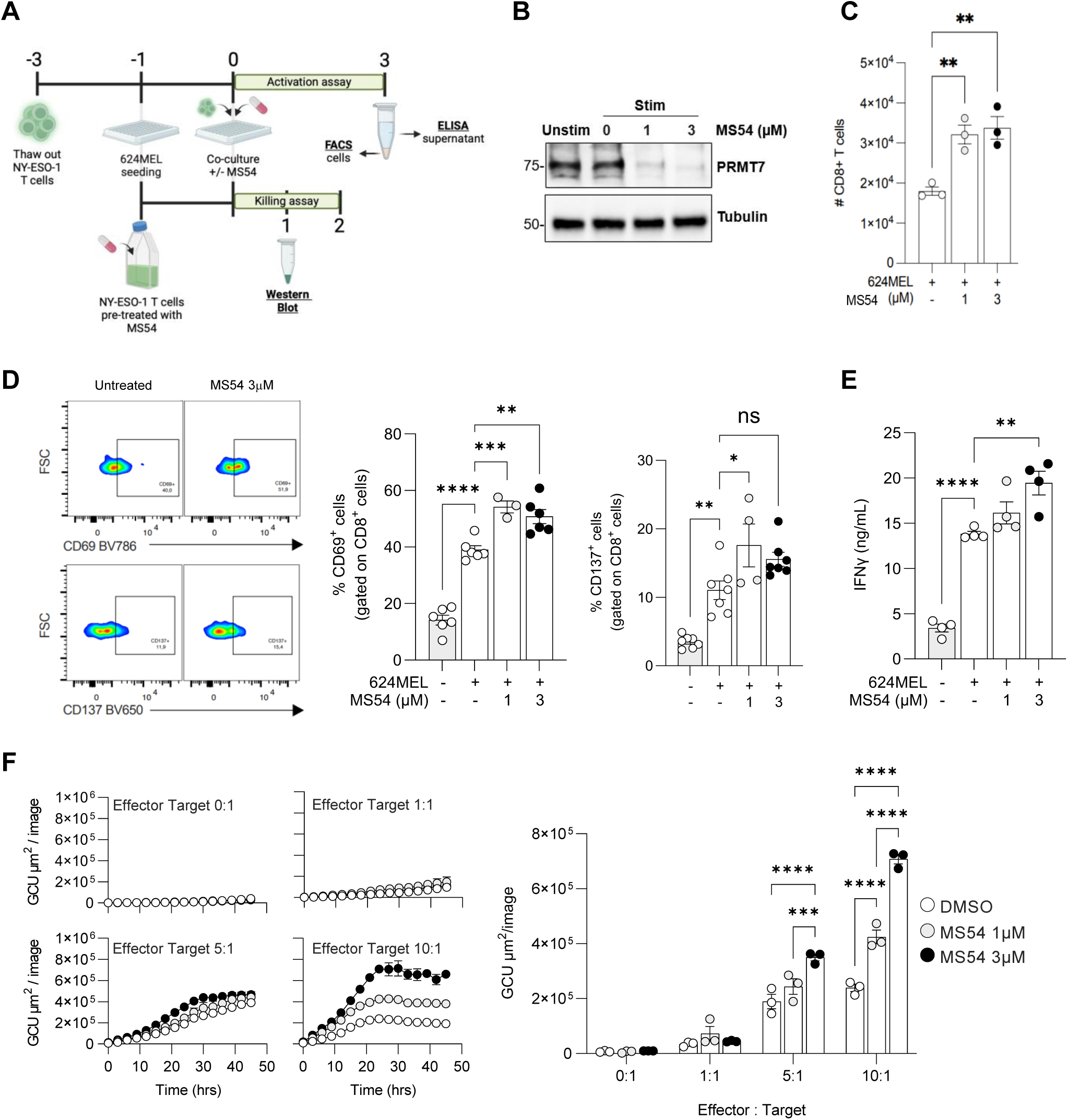
MS54 enhances human CD8^+^ T cell effector functions. **A.** Schema representing experimental procedures, timeline (in days) and readouts of the NY-ESO-1 recognition assay. HLA-matched antigen recognition assay composed of primary human T cells modified to express a transgenic TCR specific for NY-ESO-1 were co-cultured with HLA-matched 624mel human melanoma cells endogenously expressing NY-ESO-1. **B.** Western blot showing PRMT7 degradation in NY-ESO-1 T cells treated with MS54 (1 or 3 µM) and either left unstimulated or stimulated with αCD3/CD28 for 72 h. Tubulin was used as the loading control. Molecular mass markers (kDa) are indicated on the left. A representative image is shown. **C.** NY-ESO/624mel cultures were exposed to MS54 (1 or 3 µM) for 48 hours and the absolute count of CD8^+^ T cells was measured by flow cytometry. Means ± SEM are shown. Statistical significance was determined by 1-way ANOVA (\*\**p*<0.01). **D.** NY-ESO/624mel cultures were exposed to MS54 (3µM) for 48 hours and CD69 and CD137 activation markers on CD8^+^ T cells were assessed by flow cytometry. Results show representative FACS plot and pooled data from 2 independent experiments. **E.** Absolute IFN-γ levels were measured by ELISA in the supernatant (n=4). **F.** Killing assay of 624mel cells exposed to equal or 5- or 10-times more NY-ESO-1 specific T cells pre-treated with MS54 (1 or 3µM) measured by caspase 3/7 activity (fluorescence) with Incucyte® Caspase 3/7 dye for apoptosis. Results show accumulation of fluorescence over time and mean fluorescence intensity at 24h after start of the assay. Means ± SEM are shown; \**p*<0.05, \*\**p*<0.01, \*\*\**p*<0.001, \*\*\*\**p*<0.0001 by 1-way **(D-E)** or 2-way ANOVA **(F)** with multiple comparisons and Bonferroni’s correction.

## DISCUSSION

In this study, we demonstrate that T cell specific deletion of Prmt7 using CD4-Cre transgenic mice did not affect thymic T cell development but selectively increased the number of CD8⁺ T cells in the periphery, while leaving the CD4⁺ T cell population unaffected. Loss of Prmt7 enhanced CD8⁺ T cell expansion and cytokine production resulting in increased effector functions such as tumor cytotoxic killing. Transcriptomic analysis of Prmt7-deficient CD8⁺ T cells revealed transcriptional reprogramming including activation of the NF-κB signaling pathway. We developed MS54, a potent and selective PROTAC degrader of PRMT7 that is membrane-permeable and enhances the immunomodulatory activity of CD8^+^ T cells. In murine T cells, MS54 enhanced their effector function, with increased activation markers including CD25, CD69, and IFNγ expression. MS54-treated OT-I CTLs improved their *in vivo* antitumor efficacy, and this effect was increased when combined with anti-PD-L1 immunotherapy. We extend these findings to human CTLs using a validated HLA-matched antigen recognition system (Allard et al, 2025). PRMT7 loss in NY-ESO-1 TCR-engineered T cells enhanced the presence of activation markers CD69, CD137, and Ki-67, reflecting robust proliferation and effector killing of melanoma. CD137, a co-stimulatory receptor upregulated upon T cell activation, promotes CD8⁺ T cell survival and function and serves as a reliable marker of tumor-reactive cells (Stagg *et al*, 2011; Ye *et al*, 2014). Its engagement enhances proliferation, cytotoxicity and antitumor immunity in preclinical models (Khushalani *et al*, 2024; Murillo *et al*, 2008), consistent with the improved tumor killing and IFNγ secretion observed in our co-culture. In sum, our findings show that PRMT7 is a restriction factor for CD8^+^ T cells and that MS54 is a promising therapeutic to increase CD8^+^ T cell efficacy and persistence for improved adoptive T cell therapies.

The NF-κB signaling pathway plays a pivotal role in the activation, differentiation, and survival of T cells, especially CD8⁺ T cells (Gerondakis *et al*, 2014). Upon TCR engagement, NF-κB, together with AP-1 and NFAT, orchestrate transcriptional programs controlling cytokine production, proliferation, and cytotoxic effector function (Gerstberger *et al*, 2014; Ziegler, 2006). Lineage-specific studies in mice show that inhibition of NF-κB or deletion of IKKβ or NEMO dramatically reduces single-positive CD8⁺ thymocytes and peripheral CD8⁺ T cells, whereas CD4⁺ populations are comparatively less affected (Courtois & Gilmore, 2006; Hettmann & Leiden, 2000; Mora *et al*, 1999; Schmidt-Supprian *et al*, 2003). Similarly, loss of NF-κB subunits including RelA, c-Rel, or NF-κB1 impairs CD8⁺ effector differentiation and survival (Gerondakis *et al*., 2014). In our study, PRMT7 deficiency selectively enhanced NF-κB signaling in CD8^+^ T cells, upregulating genes involved in survival (Bcl family) and effector differentiation (e.g., IFNγ). This supports prior evidence that NF-κB activity is particularly indispensable for CD8⁺ lineage specification and memory maintenance (Jimi *et al*, 2008). PRMT7 may thus act as a modulatory brake on NF-κB driven transcriptional program in CD8⁺ T cells. Importantly, this controlled and transient activation of NF-κB may have translational relevance, as fine-tuning NF-κB activity could enhance CD8⁺ T cell responses and improve the efficacy of adoptive or checkpoint-based immunotherapies (Eluard *et al*, 2020; Garris *et al*, 2018). Unlike well-established negative regulators of NF-κB in CD8⁺ T cells, such asA20, CYLD, and IκBα (Dabbah-Krancher *et al*, 2024; Liu *et al*, 2017; Yin *et al*, 2022), loss of PRMT7 appears to provide a physiologically appropriate increase in NF-κB activity to enhance CD8⁺ T cells effector function. By contrast, constant NF-κB activation, for example through a constitutively active IKK mutant, triggers negative selection and apoptosis in CD8⁺ T cells (Jimi *et al*., 2008; Krishna *et al*, 2012) highlighting the importance of balanced NF-κB modulation. Interestingly, ChIP-Seq analysis shows that p65 binds the PRMT7 promoter, suggesting that NF-κB transcriptionally regulates PRMT7 expression (Gunes Gunsel *et al*, 2022). This establishes a bidirectional feedback loop where the NF-κB pathway promotes PRMT7 expression, and PRMT7 regulates p65 nuclear translocation, fine-tuning NF-κB-dependent immune responses. This preferential impact on CD8⁺ T cells may also reflect their higher intrinsic Prmt7 expression (∼4-fold higher than CD4⁺ T cells, Figure 1F), implying that PRMT7 is more functionally engaged in the CD8⁺ T cells lineage.

We observe that PRMT7 functions as a co-accessory NF-κB restriction factor that prevents p65 nuclear translocation. This contrasts with other co-accessory proteins that enhance and are required for proper NF-κB activation, including RNA-binding proteins such as RPS3 and Sam68 (Fu *et al*, 2013; Wan *et al*, 2007), as well as CARM1 (Covic et al, 2005), PRMT1 (Hassa et al, 2008), PRMT5 (Wei et al, 2013) and PRMT6 (Di Lorenzo et al, 2014). As the enzymatic activity of PRMT7 was not required to attenuate the NF-κB pathway (Figures 2G and 2H), this implied that PRMT7 has a non-catalytic function in this pathway by associating with p65. Further, we investigated whether p65 was methylated *in vitro* and *in vivo* at known RxR motifs by PRMT7 (Feng *et al*., 2013). We were unable to methylate recombinant p65 *in vitro* using recombinant PRMT7 where H2B served as a positive control (not shown).

Additionally, immunoprecipitated endogenous or epitope tagged PRMT7 did not react with anti-MMA antibodies nor were we able to detect MMA methylation on p65 in cells using trypsin or chymotrypsin followed by mass spectrometry (data not shown).

Preclinical studies have identified strategies to improve adoptive cell therapies, such as p38 kinase inhibition (Gurusamy *et al*, 2020), targeting TET2 (Dimitri *et al*, 2024), DNMT3A (Prinzing *et al*, 2021), and HDACs (Zhu *et al*, 2024), which preserve memory phenotypes and prevent dysfunction during chronic stimulation. Additional factors such as viral transduction optimization (Amiot *et al*, 1996) and spatial positioning and interactions of CD8^+^ with CD4^+^ T cells may dictate anti-tumor responses in the context of adoptive T cell therapy (Espinosa-Carrasco *et al*, 2024). As metabolic checkpoints such as AMPK and mTOR pathways also influence NF-κB mediated effector programs in CD8^+^ T cells (Chi, 2012; Rao *et al*, 2015), this suggests that combining PRMT7-targeted degradation with metabolic optimization (Blagih *et al*, 2015; Dahabieh *et al*, 2025; Ma *et al*, 2025; Roy *et al*, 2020; Sheldon *et al*, 2021), may represent a promising avenue for optimal CD8^+^ T cell expansion.

In conclusion, our study positions PRMT7 as a druggable restriction factor limiting CD8⁺ T cell effector functions. Treatment with MS54, a selective PROTAC degrader of PRMT7, effectively boosts CD8⁺ T cell adoptive cell transfer for effective cancer immunotherapy.

## ACKNOWLEDGEMENTS

We thank Dr. Steven Clarke for helpful discussions. We thank Dr. Oscar Villarreal for expert bioinformatic analysis. We are grateful to Dr. Su Jie for performing the mouse i.v. injections. We thank Marie-Lyne Fillion for expert assistance with microscopy and Christian Young at the Lady Davis Institute for support with flow cytometry. N.S. was a recipient of a PDF award from the Fonds de recherche du Québec. This work is funded by Canadian Institutes of Health Canada (CIHR) grant PJT-198035 to S.R. J.J. acknowledges the support by an endowed professorship by the Icahn School of Medicine at Mount Sinai.

## MATERIAL & METHODS

### Generation of *Prmt7^FL/FL^*; CD4-Cre mice (*Prmt7* cKO mice)

*Prmt7^FL/FL^* mice were generated by disrupting exon 4 of the *Prmt7* gene using standard LoxP gene targeting technology (Blanc *et al*., 2016). *Prmt7*^FL/FL^ mice were crossed with transgenic CD4-Cre mice (B6.Cg-Tg(Cd4-cre)1Cwi/BfluJ; JAX#022071), to generate Prmt7*^FL/FL^*; CD4-Cre (*Prmt7* cKO). These mice were bred and housed at the LDI-animal quarter; they were healthy with no signs of being immunodeficient. For ACT experiments, OT-I transgenic mice (C57BL/6-Tg (TcraTcrb)1100Mjb/J; JAX#:003831) and C57BL/6J mice (JAX#000664) were used.

### Animal Experimentation

All mouse procedures were performed in accordance with McGill University guidelines, which are set by the Canadian Council on Animal Care. Seven to twelve old wild type female mice were subcutaneously injected with 1×10^6^ cells/100μl into the right flank on day 0. On day 3, 6, 9 and 12, mice were treated with 100μl of monoclonal anti-PD-L1 (anti-mouse B7-H1, clone: 10F.9G2, Cat #BE0101 InVivoMAb). Measurements were taken manually with a caliper by collecting the longest dimension (length) and the longest perpendicular dimension (width). We estimated the tumor volume with the formula: (L×W2)/2. CO_2_ inhalation was used to euthanize mice for tumor collection. For the *in vivo* injection of NF-κB inhibitors: *Prmt7*^FL/FL^ and Prmt7 cKO 7 to 12 weeks old mice were subcutaneously injected with 1×10^6^ cells/100μl of B16.F10 melanoma cells into the right flank on day 0 and then i.p. injected with either BAY 11-7082 (MedChem Express; # HY-13453 at 5mg/kg) or Bortezomib (LC-Labs; #B-1408 at 1mg/kg); 3 times per week for 3 weeks. Tumor size and overall survival were measured and calculated as described above.

### Cell lines and transfections

B16.F10 murine cell line was kindly provided by Dr. Michael Pollak (McGill University, Montreal). These cells were subjected to CRISPR/Cas9-Mediated knockout of PRMT7 as described (Srour *et al*., 2022). 624mel human cell line (RRID: CVCL_8054) was provided by Dr. John Stagg (CRCHUM, Montreal). Cells were maintained in Dulbecco’s modifies Eagle’s medium (HyClone), supplemented with 10% Fetal Bovine Serum (FBS: HyClone), 1% penicillin/streptomycin (Multicell) and 1% sodium pyruvate (Multicell) in a 5% CO2 incubator at 37 °C. NY-ESO-1–specific T cells were thawed out 5 days prior to the coculture with 624mel and maintained in RPMI with 10% human serum (AB Gemini, cat. #100-512), 1X GlutaMAX, 1X HEPES, antibiotics, and 300 IU/mL of IL-2 (Proleukin obtained at the CHUM pharmacy). For coculture assays, 0.05 × 10^6^ cells per well of 624mel were seeded in a 96-well plate, and NY-ESO-1–specific T cells were added at a 1:1 ratio. Cells were incubated at 37°C harvested after 48h or 72h for further analysis (Flow cytometry and Western Blot). Supernatant was stored at −20°C for further analysis by ELISA. NY-ESO-1–specific T cells were obtained from Dr. Simon Turcotte (CRCHUM, Montreal). For vector transfections, B16.F10 cells were transfected using Lipofectamine 3000 (Invitrogen), according to the manufacturer’s instruction.

### Flow Cytometry

For immune cell phenotyping, T cells freshly isolated from mice (splenocytes and thymocytes) were blocked with anti-mouse CD16/CD32 (Fc block; BD Biosciences) for 15 min and stained with indicated fluorescence-conjugated antibodies for 30 min as well as a Live/Dead discrimination dye (BD Biosciences).

For tumor-infiltrating lymphocytes (TIL) analysis, primary tumors were collected at end point, weighed, mechanically diced, and digested in RPMI + 1 mg/ml in the presence of collagenase P (2 mg/ml, Sigma-Aldrich) and DNase I (50 μg/ml, Sigma-Aldrich) for 30 min at 37°C. Next, cells were pipetted into a single-cell suspension, blocked with anti-mouse CD16/CD32 antibody (Fc block; BD Biosciences) for 15 min and stained with fluorescence-conjugated antibodies for 30 min and then acquired with a BD LSRFortessa flow cytometer and analyzed using FlowJo v10.8.0 software. Gating strategy was based on fluorescence minus one (FMO) control. Beads were used for spectral overlap compensation.

### Real-time imaging of T cell-mediated killing assay using the Incucyte system

For killing assay using murine B16.F10 melanoma cells, cancer cells were seeded in 96-well flat-bottom plates at 2,000 cells/well in 100 µL complete medium and incubated overnight to reach ∼20% confluency. The following day, cells were incubated with Caspase-3/7 Green apoptosis dye (5 µM; Essen Bioscience #4440) and subjected to the indicated experimental conditions. T cells were freshly isolated from spleens, pre-activated with anti-CD3/anti-CD28 beads and treated with vehicle or MS54 (3µM) for 24 h prior the co-culture. They were then added to the wells at the indicated effector-to-target (E: T) of 1:5 in 100 µL, resulting in a final assay volume of 200 µL per well.

For killing assay of the human HLA-matched cells, 3×10^3^ 624mel cells were seeded per well of a 96-well plate the day before the assay. NY-ESO-1 specific T cells were pretreated with MS54 (1-3µM) or DMSO for 24 h prior to co-culturing. On the day of the assay, cells were treated with Caspase-3/7 dye as described previously and NY-ESO-1 T cells were added at effector-to-target (E:T) ratios of 1:1, 5:1, or 10:1. Plates were equilibrated at room temperature for 30 min and then placed in the Incucyte® Live-Cell Analysis System. Images were acquired every 30 min or 2 h using phase contrast and green fluorescence channels with a 10x objective. Apoptotic cell death was quantified over time based on Caspase-3/7 fluorescence using the Incucyte® analysis software.

### Protein extracts and immunoblot analysis

Whole lysates from B16.F10 melanoma cells and from murine and human T cells were prepared in 2x Laemmli buffer and boiled at 100°C. Protein extracts were resolved by SDS-PAGE, transferred to nitrocellulose membranes using an immunoblot TurboTransfer system (Bio-Rad), blocked for 1h at room temperature in TBS-T 5% milk and incubated with corresponding primary antibodies, followed by incubation of secondary antibodies conjugated to horseradish peroxidase (Sigma Aldrich). Immunoblot signals were detected using chemiluminescence (Perkin Elmer).

### Immunoprecipitation

For immunoprecipitations, cells were lysed in lysis buffer (1% Triton, 150 mM NaCl, 20 mM Tris-HCl pH 8.0, 100 mM sodium vanadate, 0.01% phenylmethanesulfonyl fluoride and protease inhibitors). Next, the lysates were cleared, 50 μl of the lysates was collected as input and 500 μl was used for immunoprecipitation with 1 μg of anti-p65 antibody (2h on ice) or mouse IgG and protein A agarose beads (40 μl/50% slurry) for endogenous p65 or incubated with anti-Flag affinity beads (40 μl /50% slurry) at 4°C for 1 h under constant rotation for transfected Flag-tagged p65. The beads were then washed 4 times with lysis buffer, boiled with 40 μl of 2× Laemmli buffer and immunoblotted.

### Nuclear/Cytoplasmic Fractionation

B16.F10 melanoma cells were treated with BAY11-7082 (10 μM) or vehicle for 6 h. Nuclear and cytoplasmic fractions were isolated using the Abcam Nuclear/Cytoplasmic fractionation Kit (ab113474), following the manufacturer’s instructions. Briefly, cells were washed with ice-cold phosphate-buffered saline (PBS) and harvested using trypsin. After centrifugation, the cell pellet was resuspended in a hypotonic buffer containing protease inhibitors. Following incubation on ice, detergent was added to disrupt cell membranes, and nuclei were pelleted by centrifugation. The supernatant, containing the cytoplasmic fraction, was collected, and the nuclear pellet was resuspended in extraction buffer to isolate the nuclear fraction. Fractions were analyzed by SDS-PAGE and immunoblotting with an anti-p65 antibody. To ensure the purity of the fractions, Tubulin was used as a positive control for the cytoplasmic marker and a negative control for the nuclear fraction.

### Dual-luciferase reporter assay

sgCTL and sgPRMT7 B16.F10 melanoma cells were seeded in 24-well plates and then transiently co-transfected in triplicate with the NF-κB promoter-Luc reporter, alone or with the indicated expression vectors (FLAG-p65, GFP-PRMT7^WT^ or GFP-PRMT7^DEAD^). The p65 plasmid and NF-κB promoter-luc reporter was provided by Dr. Rongtuan Lin, (McGill University) (Belgnaoui *et al*, 2012). The GFP-tagged PRMT7-WT and the catalytically inactive mutant PRMT7^DEAD^ plasmids were provided by Dr. Steven Clarke (UCLA) (Feng *et al*., 2013). The Renilla pRL-TK plasmid was used as an internal control, and the total amounts of DNA were kept constant by supplementation with an empty vector (pcDNA3.1). After 24 h, cell lysates were harvested, and the relative luciferase units (RLUs) were measured using the dual-luciferase assay according to the manufacturer’s instructions (Promega #E1910). RLUs from firefly luciferase signal were normalized by RLUs from Renilla signal.

### Immunofluorescence

Immunofluorescence staining was performed using a similar protocol for both B16.F10 melanoma cells and human T cells (CCRF CEM). B16.F10 cells were cultured on glass coverslips, while human T cells were prepared by cytospin (100–200 μL cell suspension at ∼5×10^5^ cells) onto glass slides using a cytocentrifuge (1000xg, 5 min). Cells were washed twice with PBS and fixed for 10 min with 4% paraformaldehyde (PFA). After three washes, cells were permeabilized for 5 min with 0.5% Triton X-100 in PBS and blocked overnight in PBS containing 10% FBS and 0.2% Triton X-100. Samples were incubated with anti-p65 antibody (CST #8242S), diluted (1/200) in PBS containing 5% FBS for 1 h at room temperature, washed three times, and then incubated with the corresponding fluorescent secondary antibody for 30 min in PBS with 5% FBS. After rinsing, coverslips were mounted using Immuno-Mount (Thermo Scientific) mounting medium containing 1 μg/mL DAPI. For imaging, B16.F10 samples were acquired on a Zeiss Axio Imager M1 fluorescence microscope and analyzed using ImageJ software, whereas human T cells (CCRF-CEM) were imaged on a Nikon AXR point laser scanning confocal microscope and analyzed using GA3 (General Analysis 3) in NIS-Elements software. Magnification 60x and numerical aperture of the objective:1.42 NA.

### RT-qPCR

Total RNA from cells were isolated with TRIzol (Invitrogen) according to manufacturer’s instruction and quantified by NanoDrop. After digestion with DNase I (Promega), 1 μg of total RNA was converted to cDNAs using M-MLV reverse transcriptase (Promega). Real-time quantitative PCRs were performed using PowerUp SYBR Mastermix (Life Technologies #A25742) on 7500 Fast Real-Time PCR System (Applied Biosystem). Results were normalized as described in the figure legends using the ΔΔct method. Primers used in this study are outlined in Table 1.

### Human T cell-antigen recognition assay

624mel (RRID: CVCL_8054) cells were routinely tested for mycoplasma (MycoAlert, Lonza #LT07-318) and cultured in DMEM containing 10% FBS. NY-ESO-1 specific T cells were expanded *in vitro* according to a previously described standard rapid expansion protocol (Berry *et al*, 2025). Both 624mel and NY-ESO-1 specific T cells were obtained from Dr. Simon Turcotte (CR-CHUM, Montreal, QC, CA). NY-ESO-1 specific T cells were thawed out 3 days prior to the co-culture with 624mel and maintained in RPMI with 10% human serum (AB Gemini #100-512), 1X Glutamax, 1X HEPES, antibiotics and 300 IU/mL of IL-2 (Proleukine obtained at the CHUM pharmacy). For activation assays, 0.05×10^6^ cells per well of 624mel were seeded in a 96-well plate, and NY-ESO-1-specific T cells were added in a IL2-free medium at a 2:1 ratio. Cells were co-cultured at 37°C for 3 days, labeled for CD45, CD4, CD8, CD69, CD137 and Ki-67, and viability, activation and absolute cell counts were analyzed by flow cytometry. In parallel, supernatants were collected and analyzed for IFNγ levels by ELISA according to a previously described protocol (Allard et al, 2025). For PRMT7 degradation measurements by western blot, NY-ESO-1 specific T cells were pre-treated for 24 h prior to co-culture with 624mel for another 24 h. Whole protein extracts were then analyzed by western blot as described previously. For killing assays, 3×10^3^ 624mel cells were seeded per well of a 96-well plate the day before the assay. NY-ESO-1 specific T cells were pretreated with MS54 compound (1-3µM) or DMSO for 24 h prior the co-culture and T-cell killing was measured as described previously.

### Measurements of IFNγ by ELISA

Supernatants from cocultures of NY-ESO-1-specific T cells and 624mel cells were collected after 3 days and analyzed for IFNγ levels by ELISA. Briefly, 96-well Nunc MaxiSorp plates were coated overnight at 4°C with 1 μg/mL of primary anti–IFNγ antibody (clone 2G1) in 100 μL PBS per well. Plates were washed with 0.2% Tween20 PBS, blocked 60 min with 1% BSA PBS and incubated for 90 min with diluted supernatant and biotinylated IFNγ antibody (clone B133.5; Thermo Fisher Scientific #M701B, RRID: AB_223580). To reveal, plates were washed, incubated with HRP-conjugated streptavidin (Fitzgerald #55R-S103PHRP) for 30 min, washed again, incubated with enhanced K-blue TMB substrate (Neogen #308177) for 5 min and absorbance was measured on a Varioskan™ (Thermo Fisher Scientific) instrument at 450 nm OD values were reported on a standard curve made with recombinant IFNγ (Thermo Fisher Scientific, cat. #EN-RIFNG50) to calculate absolute levels.

### RNA sequencing and data analysis

RNA samples were purified using GenElute™ Mammalian Total RNA Miniprep Kit (RTN70, Sigma Aldrich). Total RNA was assessed for quality using an Agilent Tapestation 4200, and RNA sequencing libraries were generated using TruSeq Stranded mRNA Sample Prep Kit. Samples were processed following manufacturer’s instructions, starting with 50 ng of RNA and modifying RNA shear time to 5 min. Resulting libraries were multiplexed and sequenced with 100 base pair (bp) to a depth of approximately 30 million reads per sample. Samples were demultiplexed using bcl2fastq v2.20 Conversion Software (Illumina, San Diego, CA). Reads were mapped to the Genome Reference Consortium Mouse Build 38 patch release 6 (mm10/GRCm38.p6: primary assembly) (Frankish *et al*, 2019) using STAR v2.4 (Dobin *et al*, 2013).

### Gene expression analysis

Expression levels were estimated using HOMER V4.10 (Heinz *et al*, 2010). Afterwards, we employed DESeq2 (Love *et al*, 2014) to normalize the raw counts as rlog variance stabilized values, as well as to perform the differential expression analysis as previously described (Darbelli *et al*, 2017). For the volcano plot, genes were considered differentially expressed if they had an adjusted p value <0.05, a base mean higher than 100 and an absolute fold-change greater than 2. For the heat map, genes were considered differentially expressed if the samples with the highest and lowest expression are more than 2-fold different and one of the samples has 25 normalized reads (as in the HOMER tutorial).

### Gene ontology

GO term enrichment analysis of the differentially expressed genes was performed through one or more of the following: STRING v11.0, GSEA V3.0, Enrichr, DAVID, and IPA. The list of differentially expressed genes was compared to a background of expressed genes, consisting of all expressed genes in the complete dataset as defined by all genes with the DESeq2 base mean higher than the first expression quartile. For differentially expressed genes, upregulated and downregulated genes with a base mean higher than 100 and an absolute fold-change greater than 2 were used for the analysis.

### Chemistry General Procedures

All chemical reagents were purchased from commercial vendors and used without further purification. Flash column chromatography was performed on a Teledyne ISCO CombiFlash Rf^+^ instrument equipped with a 220/254/280 nm wavelength UV detector and a fraction collector. Reversed-phase column chromatography was conducted on HP C18 RediSep Rf columns to purify the polar compounds. All final compounds were purified with preparative high-performance liquid chromatography (HPLC) on an Agilent Prep1200 series with the UV detector set to 220 and 254 nm at a flow rate of 40 mL/min. Samples were injected onto a Phenomenex Luna 750× 30 mm, 5 *μ*m C18 column, and the gradient was set to 10% acetonitrile in H_2_O containing 0.1% TFA progressing to 100% acetonitrile. The purity of all compounds was determined by an Agilent 1200 series system with DAD detector and a 2.1 mm x 50 mm InfinityLab Poroshell 120 EC-C18 column for chromatography. Samples (0.5 µL) were injected onto a C18 column at room temperature with the flow rate of 0.6 mL/min. Chromatography was performed with the solvent as follows: water containing 0.1% formic acid and 3% acetonitrile was designated as Solvent A while acetonitrile containing 0.1% formic acid was designated as solvent B. The linear gradient was set such that 5% B was used from 0 – 0.5 min, 5 – 100% B from 0.5 – 2.8 min, 100% B from 2.8 – 4.2 min, 100 – 5% B from 4.2 – 4.3 min, and 5% B from 4.3 – 5 min. High-resolution mass spectra (HRMS) data was acquired in positive ion mode using an Agilent 6230 TOF with an electrospray ionization (ESI) source. Nuclear magnetic resonance (NMR) spectra were acquired on either Bruker DRX 400 MHz for proton (^1^H NMR) and 101 MHz for carbon (^13^CNMR). Chemical shifts for all compounds are reported in parts per million (ppm, δ). The format of the chemical shift was reported as follows: chemical shift, multiplicity (s = singlet, d = doublet, t =triplet, q = quartet, m = multiplet), coupling constant (J values in Hz), and integration. All final compounds had >95% purity using the HPLC methods described above. For the synthesis route of MS54 see Supplementary Methods (PRMT7^PRO^ synthesis).

### Statistical analysis and data representation

All experiments were repeated at least two to three times, except as specified otherwise. All data are presented as mean ± standard error of the mean (SEM). GraphPad Prism software (version10.3.1) was used to generate plots and additional statistical analysis. Significance of comparison between two groups was assessed either by the unpaired or paired Student-t test. The use of the specific tests as well as the number of animals and experimental replicates has been reported in each figure legend. Statistically significant results were defined as follows: *\*p <0.05; **p <0.01; ***p <0.001; ****p <0.0001*. Statistical analysis for RNA-seq was performed with DESseq2 for gene expression.

## Data availability

The RNA-seq data that support the findings of this study have been deposited in the NCBI Gene Expression Omnibus and are accessible through the accession number GSE309747.

## SUPPLEMENTAL FIGURE LEGENDS

**Figure S1. *Prmt7* cKO mice has no effect on Thymic T cell development (Related to Figure 1).**

**A.** Average RNA expression of PRMT7 across various tissues.

**B.** PRMT7 expression across human immune cell types (GSE107011). Boxplots show TPM-normalized expression of PRMT7 across different immune cell populations.

**C.** PRMT7 expression in naive T cell subsets. PRMT7 mRNA expression is shown for T4 naïve (blue) and T8 naïve (green) samples. Values are DESeq2-normalized counts (size-factor normalization), and each vertical bar represents an individual sample. The y-axis shows DESeq2-normalized counts (linear scale).

**D.** Representative flow cytometry plots showing CD62L and CD44 expression in splenocytes isolated from control and *Prmt7* cKO mice gated on CD4^+^ T cells.

**E.** Quantification graphs from (**D)** showing the percentages of CD62L^+^CD44^-^ (naïve), CD62L^+^CD44^+^ (central memory) and CD62L^-^CD44^+^ (effector memory) T cell subsets gated on CD4^+^ T cells, in *Prmt7^FL/FL^*control (black dots) and *Prmt7* cKO (red dots) mice (n=3).

**F.** Representative flow cytometry plots showing CD45 and NK1.1 expression in splenocytes isolated from control and *Prmt7* cKO mice.

**G.** Percentage of iNKT cells gated on total CD45^+^ cells in *Prmt7^FL/FL^* control (black dots) and *Prmt7* cKO (red dots) (n=6). **(D-G).** Data are representative of two to three independent experiments. Mean ± SEM is shown. Statistical significance was determined using unpaired Student’s *t*-test (\*\**p* <0.01; *ns*: not significant).

**Figure S2. Transcriptomic profiling of *Prmt7* cKO T cells (Related to Figure 3).**

**A.** Principal component analysis (PCA) of transcriptomic profiles in CD8^+^ T cells isolated from *Prmt7^FL/FL^* control and *Prmt7* cKO mice (n=3/group). PCA was performed on DESeq2- normalized RNA-seq counts. Each point represents an independent biological replicate of PRMT7CTL (pink) or PRMT7KO (blue) samples. PC1 and PC2 account for 50% and 22% of the variance, respectively, and clearly separate PRMT7 knockout from control groups.

**B.** Gene Ontology (GO): Pathways enriched among the top differentially expressed genes between PRMT7 CTL and KO samples (q-value < 0.05; LFC < –3 for downregulated, LFC > 3 for upregulated).

**C.** Enrichment dot plot showing the top 50 transcription factors ranked by mapped peaks ratio, defined by the percentage of promoters of differentially expressed genes (DEGs) bound by each transcription factor.

**D.** Genes with the most differential target genes for RelA were identified using decoupleR. TFs regulating RelA are outlined in red and blue, where red indicates activation and blue indicates repression. This panel highlights the key TFs that regulate RelA.

**E.** Heatmap showing expression patterns of MAPK pathway–related genes in PRMT7 CTL and KO samples.

**F.** Predicted full-length structures of PRMT7 and RelA generated by AlphaFold 3 with the corresponding predicted alignment error (PAE) plot. Predictions were obtained from the AlphaFold server. PyMOL-processed representation of the AlphaFold 3 structural models. PRMT7 is shown in green, with its first 100 residues highlighted in yellow. RelA is shown in blue, with the dimerization domain highlighted in cyan. A zoomed-in image highlights the interface where the β-sheet in the PRMT7 N-terminus aligns with the β-sheet in the p65 dimerization domain. Images were generated using PyMOL v3.1.4.1.

**Figure S3. PRMT7 affects NF-κB activation by retaining p65 in the cytoplasm (Related to Figure 3).**

**A.** Representative immunofluorescent images of sgCTL and sgPRMT7 B16.F10 melanoma cells treated with TNF-α (50 ng/mL) for 20 min and stained with anti-p65 antibody at 40x magnification. Nuclei were visualized with DAPI (4’,6-diamidino-2-phenylindole, blue) using a Zeiss confocal microscope. Scale bar = 10 µm.

**B.** Representative immunofluorescence images of B16.F10 cells transfected with GFP-PRMT7 and stained for PRMT7 (green) and p65 (red) at 40× magnification. Nuclei were visualized with DAPI (4’,6-diamidino-2-phenylindole, blue) using a Zeiss confocal microscope. Scale bar = 10 µm. Three different fields are shown (#1, #2, #3).

**C.** Quantification of nuclear and cytoplasmic intensities in GFP⁻ (non-transfected) and GFP⁺ (transfected) cells observed in **(B)**. Fluorescence intensities were measured in ImageJ by drawing nuclear and cytoplasmic ROIs on single-cell planes, with background subtraction applied to all measurements.

**D.** Western blot of Cytoplasmic (C) and Nuclear (N) fractions showing Prmt7 expression from T cells isolated from *Prmt7*^FL/FL^ control mice. Tubulin and Histone (H3) were used as a loading control for the cytoplasmic and nuclear fractions respectively. Molecular mass markers (kDa) are indicated on the left. Two independent experiments are shown (#1, #2).

**Figure S4. Lymphoid and Myeloid TIL profiles (Related to Figure 4).**

**A.** B16.F10 melanoma cells were subcutaneously injected in *Prmt7^FL/FL^* control and *Prmt7* cKO mice followed by intraperitoneal injection of either BAY 11-7082 (5mg/kg) or Bortezomib (1mg/kg) in *Prmt7* cKO mice. At endpoint, tumors were digested into single-cell suspensions and immune cell composition was analyzed. Quantification of lymphocyte populations, including total CD3⁺, CD4⁺, CD8⁺, regulatory T cells (Tregs), and NK cells.

**B.** Quantification of Myeloid-Derived Suppressor Cell (MDSC) populations, including Granulocytic MDSC (G-MDSC, F4/80^-^), Monocytic MDSC (M-MDSC, F4/80^+^), neutrophils, and dendritic cells. Data represent mean ± SD from two independent experiments with a minimum of 5 mice per group. Statistical significance was calculated using two-way ANOVA. (\*\**p* <0.01; \*\*\**p* <0.001; \*\*\*\**p* <0.0001; *ns*: not significant).

**Figure S5. Design and screening of putative PRMT7 PROTAC degraders (Related to Figure 5).**

**A.** Cocrystal structure of SGC8158 in complex with PRMT7 (PDB ID: 6OGN). Red dash circle indicates the solvent exposure region.

**B.** Chemicals structures of putative PRMT7 PROTAC degraders (13 compounds).

**C.** Immunoblot of PRMT7 in B16.F10 cells treated with indicated compound at 3 and 10 μM for 24 h. Tubulin was used as a loading control.

**Figure S6. MS54 treated CD8^+^ T cells have increased effector functions (Related to Figure 5).**

**A.** RT-qPCR analysis of the PRMT7 mRNA level in B16.F10 cells treated with DMSO or MS54 (10 μM) at 24 h or 48 h.

**B.** Freshly isolated T cells from control (n=6) mice were treated with vehicle or 3 μM MS54. T cells were obtained using Pan-T cell isolation kit and stimulated with αCD3/CD28 Abs for 72h. Phase contrast images show the number of colonies in T cells treated with vehicle or MS54.

**C.** Representative flow cytometry plots showing CD4 and CD8 expression in splenocytes isolated from control mice, stimulated with αCD3/CD28 Abs for 72h and treated with vehicle or 3 μM MS54. Rectangular gates identify SP CD4⁺ and SP CD8⁺ populations; percentages are shown as percent of live CD3⁺ singlets.

**D.** Quantification graphs from **(C)** showing the percentages of CD4^+^ and CD8^+^ T cell populations in vehicle-treated (black dots) and MS54-treated cells (purple dots).

**E.** Representative flow cytometry plots showing CD25 and CD69 expression on gated CD4⁺ (top) and CD8⁺ (bottom) T cells from vehicle (black) or MS54-treated cells (purple; n = 6). Quadrants indicate CD25⁺, CD69⁺, and double-positive populations.

**F.** Quantification of the percentage of CD25⁺ and CD69⁺ cells within the CD4⁺ and CD8⁺ T-cell compartments in stimulated T cells non treated (black) or treated with MS54 (purple; n = 6). Data are representative of 2 independent experiments. Means ± SEM are shown. Statistical significance was determined using unpaired Student’s *t*-test (\*\**p* <0.01; \*\*\**p* <0.001; \*\*\*\**p* <0.0001; *ns*: not significant).

**Figure S7. MS54 enhances human CD8^+^ T cell effector functions (Related to Figure 7).**

**A.** Gating strategy for CD69 and CD137 expression analysis of the NY-ESO-1 antigen recognition assay.

**B-C.** Representative FACS plot **(B)** and mean fluorescence intensity (MFI) of Ki-67 **(C)** labeling in NY-ESO-1 specific T cells co-cultured with 624mel cells. Means ± SEM are shown; \**p*<0.05, by 1-way ANOVA.

---

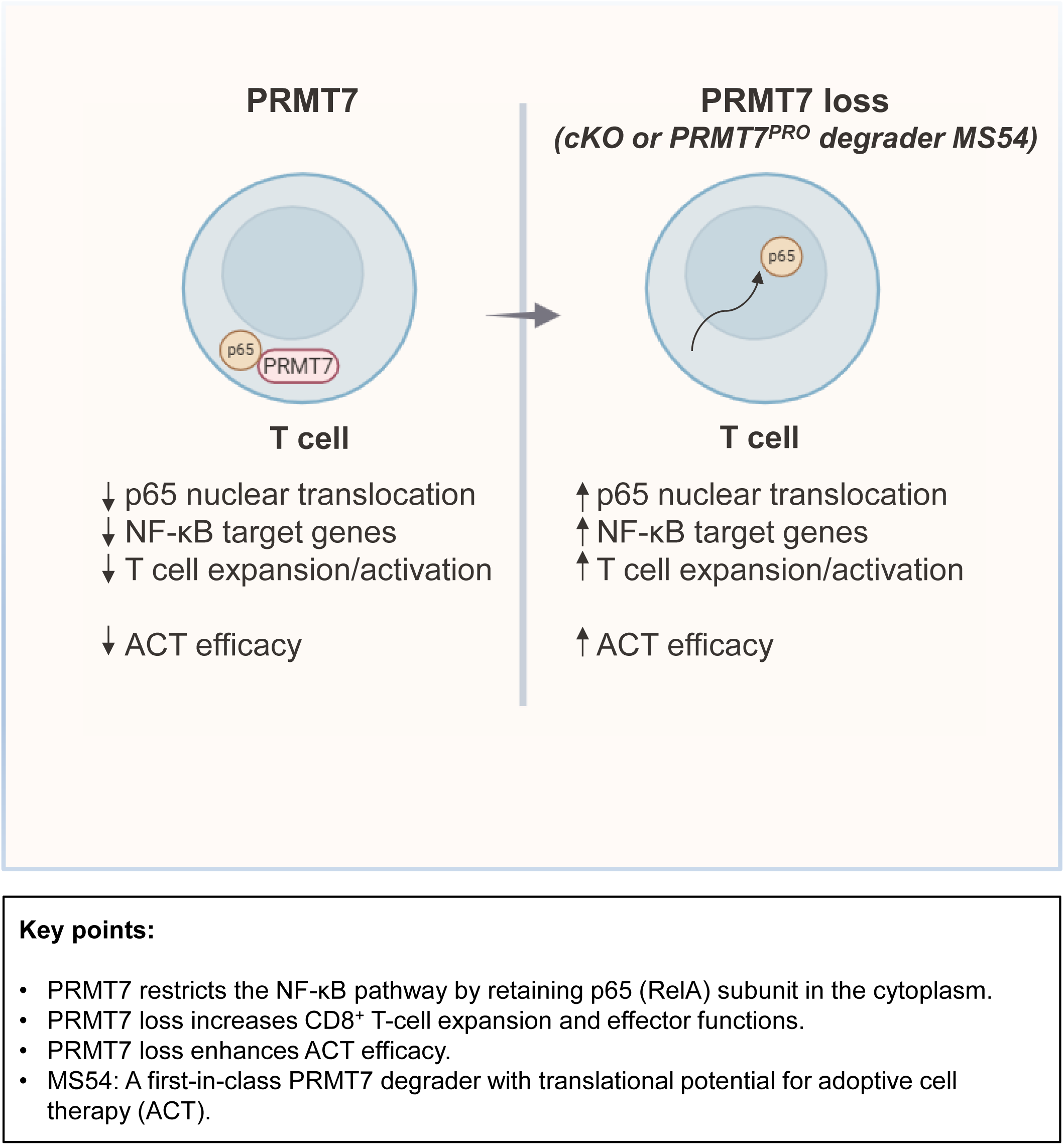

